# A molecular map of lymph node blood vascular endothelium at single cell resolution

**DOI:** 10.1101/2020.03.13.991604

**Authors:** Kevin Brulois, Anusha Rajaraman, Agata Szade, Sofia Nordling, Ania Bogoslowski, Denis Dermadi, Milladur Rahman, Helena Kiefel, Edward O’Hara, Jasper J Koning, Hiroto Kawashima, Bin Zhou, Dietmar Vestweber, Kristy Red-Horse, Reina Mebius, Ralf H. Adams, Paul Kubes, Junliang Pan, Eugene C Butcher

**Affiliations:** Laboratory of Immunology and Vascular Biology, Department of Pathology, Stanford University School of Medicine, Stanford, California, USA.; Palo Alto Veterans Institute for Research, Palo Alto, California, USA.; Calvin, Phoebe & Joan Snyder Institute for Chronic Diseases, Cumming School of Medicine, University of Calgary, Calgary, Alberta T2N 4N1, Canada.; Department of Physiology and Pharmacology, Cumming School of Medicine, University of Calgary, Calgary, Alberta T2N 4N1, Canada.; Amsterdam UMC, VU University of Amsterdam, Department of Molecular Cell Biology and Immunology, Amsterdam Infection and Immunity Institute, Amsterdam, Netherlands; Department of Biochemistry, School of Pharmacy and Pharmaceutical Sciences, Hoshi University, Tokyo, Japan.; The State Key Laboratory of Cell Biology, CAS Center for Excellence in Molecular Cell Science, Shanghai Institute of Biochemistry and Cell Biology, Chinese Academy of Sciences, 200031, China.; Department Vascular Cell Biology, Max Planck Institute for Molecular Biomedicine, Münster, Germany.; Department of Biology, Stanford University, California, USA.; Max Planck Institute for Molecular Biomedicine, Department of Tissue Morphogenesis, University of Münster, Faculty of Medicine, Münster, Germany.; The Center for Molecular Biology and Medicine, Veterans Affairs Palo Alto Health Care System, Palo Alto, California, USA

## Abstract

Blood vascular endothelial cells (BECs) control the immune response by regulating immune cell recruitment, metabolite exchange and blood flow in lymphoid tissues. However, the diversity of BEC and their origins during immune angiogenesis remain poorly understood. Here we profile transcriptomes of BEC from mouse peripheral lymph nodes and map key phenotypes to the vasculature. Our analysis identifies multiple novel subsets including a venous population whose gene signature predicts an unexpectedly selective role in myeloid cell (vs lymphocyte) recruitment to the medulla, confirmed by 2 photon videomicroscopy. We define five phenotypes of capillary lining BEC including a capillary resident regenerative population (CRP) that displays stem cell and migratory gene signatures and contributes to homeostatic BEC turnover and to vascular neogenesis after immunization. Trajectory analyses reveal retention of developmental programs along a progression of cellular phenotypes from CRP to mature venous and arterial BEC subsets. Overall, our single cell atlas provides a molecular blueprint of the lymph node blood vasculature and defines subset specialization for immune cell recruitment and vascular homeostasis.

## Introduction

The vascular endothelium lining blood vessels regulates exchange of oxygen and metabolites between the blood vascular compartment and tissues. In lymph nodes (LN), additionally, the endothelium plays essential roles in controlling immune cell access. The organization of the vasculature in LN is well characterized: Arteries entering at the hilus lead to capillary arcades in the LN cortex; capillaries link to ‘high endothelial venules’ (HEV), the post capillary venules that recruit lymphocytes from the blood^1^; and the vasculature exits the lymph node at the hilus. Upon immune challenge, lymph nodes increase in volume up to 10-fold or more within days, and the vascular endothelium expands roughly proportionally from local precursors^1,2^. Vessel expansion involves extensive proliferation of both capillary and high endothelial cells (HEC)^3^, without contribution from blood borne progenitors^4^; and an increase in vessel numbers through intussusceptive (splitting) angiogenesis^5^. However, the nature and extent of endothelial cell diversity within LN remains incompletely understood.

Single cell RNA profiling is a transformative technology for the identification of cell diversity and elucidation of developmental and physical relationships. Here we provide a survey of blood vessel endothelial cells (BEC) from mouse peripheral lymph nodes (PLN). We confirm known features of high endothelium^6,7^, identify novel endothelial subsets, uncover unexpected diversity among capillary cells and demonstrate a distinctive role of medullary veins in selective myeloid cell recruitment. We define gene signatures and transcription regulatory factors for these subsets, and map key subsets to the vasculature. We also identify a primed capillary resident regenerative population (CRP) that displays the angiogenic endothelial marker *Apln,* is enriched in cells undergoing cell division, and possesses stem cell and migratory gene signatures. Genetic lineage tracing suggests that CRP contribute to neogenesis of the blood vascular endothelium in immune angiogenesis.

## Results

### Single cell profiling of lymph node blood endothelial cells

We performed single cell RNA sequencing (scRNAseq, 10x Chromium) of sorted BEC from PLN of adult mice (**Fig. 1a**). Four cohorts were analyzed including a group of male and female Balb/c mice (PLN1) processed together and resolved *in silico* (PLN1_m and PLN1_f), an additional group of female Balb/c mice (PLN2), and one of female mice of a mixed background (PLN3) (**Supplementary Fig. 1a**). Unsupervised analyses (Methods) defined 8 BEC subsets with distinct gene expression: arterial EC (Art); 2 venous EC subsets, high endothelial cells (HEC) and non-HEC veins (Vn); and 5 capillary phenotype EC subsets, including a transitional phenotype capillary EC (TrEC) and primed EC comprising a capillary resident regenerative population (CRP) (**Fig. 1b-d**). Annotations of the arterial and venous populations were guided by known gene markers^6,8,9^. Nearest neighbor alignments, which can model developmental relationships or spatial alignments of cells, were visualized by trajectory inference using tSpace^10^, which revealed a continuum of cell phenotypes with branching of arterial and venous subsets from capillary EC populations (**Fig 1e**). Clusters extending to the termini of the arterial, CRP, and HEV branches were further separated to distinguish cells that were most distinct (distant along trajectories) from the bulk of EC (darker shading, **Fig. 1e**): these cells are enriched for genes associated with mature arterial (e.g. *Gkn3*, *Bmx*) or HEV (*Chst4*, *Glycam1*) differentiation, or in the case of CRP, for genes associated with endothelial specification during early developmental or with stem cells (e.g. *Ets2, Cxcr4, Nes*; (**Fig. 1f**). Clusters and trajectory alignments were shared by male and female mice and by the independently processed samples (**Supplementary Fig. 1b-d**). Gene expression signatures were generated for each subset using the combined scRNAseq datasets (Methods; **Supplementary Table 1**). Correlation in mean gene expression profiles of the identified subsets across the cohorts is shown in **Supplementary Fig. 1e**. Expression of marker genes depicted in **Fig. 1f** showed overall consistency across all technical replicates (**Supplementary Fig. 2**).

**Figure 1.**
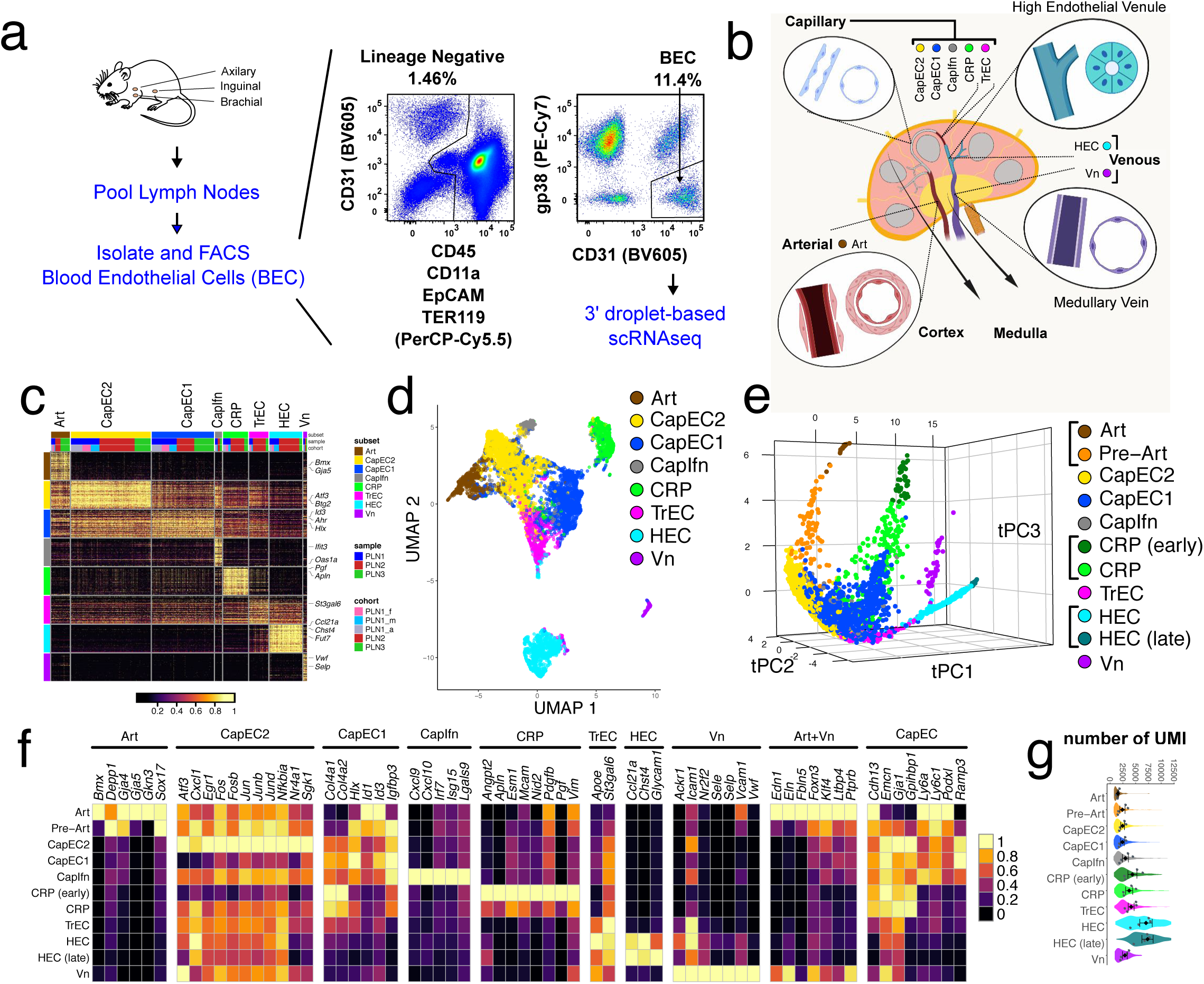
Single-cell survey of lymph node blood vessel endothelial cells. (a) Workflow schematic. Lymph nodes from adult mice are pooled and dissociated into single cells. Fluorescence activated cell sorting (FACS) is used to isolate blood endothelial cells. scRNAseq (10x Chromium) is used to profile the cells. Representative FACS plot. (b) Lymph node schematic depicting the 8 major subsets identified by scRNAseq analysis. (c) Heatmap of expression of the top 50 differentially and specifically expressed genes for each subset are shown. Subset, sample, and cohort are annotated across the top and select genes on the right. (d) UMAP plot of 8832 single cells from 3 samples (4 cohorts). Cells are segregated by type: arterial EC (pre-Art), high endothelial cells (HEC), non-HEC veins (Vn), and 5 capillary phenotype EC (CapEC1, CapEC2, capillary resident progenitors (CRP), transitional EC (TrEC), Interferon-stimulated gene-enriched CapEC (CapIfn). (e) Computationally predicted relationships visualized in PCA projection of cells aligned in trajectory space using cells from PLN1. Clusters extending to the termini of the arterial, CRP, and HEV branches were further subdivided to distinguish cells most distinct (distant along trajectories) from the bulk of EC (CRP (early), Art, HEC (late; darker shadings). Interactive rendering available: https://stanford.io/2qzJ8Hl. (f) Selected marker and signature genes for each of the indicated clusters and combinations of clusters (top). Mean expression values from 3 samples and 4 cohorts (color scale). (g) Total transcript counts (UMI; unique molecular identifiers) per cell within each subset. Violins show the UMI distribution of all cells. Mean expression values for each of the 4 independent cohorts (grey dots) and mean and standard error (SEM) of the cohort means are also plotted (black diamonds, bars).

### Characterization of the arterial and venous subsets

The arterial cluster among the profiled BEC was identified by expression of *Gja5* and *Gja4* (encoding connexin 37 and 40, respectively) and *Bmx*^11^ (**Fig. 1f)**. Consistent with prior reports describing *Gkn3* as a marker for mature arteries^8^, it is selectively expressed in mature Art, and absent in pre-Art, which lay closer to the capillary subsets in trajectory space (**Fig. 1e**). On the other hand, *Depp1* is preferentially expressed in pre-Art, consistent with a previous study showing its heterogeneous expression in arteries, including transient expression in developing arterial endothelial cells and subsequent down-regulation in mature vessels^12^ (**Fig. 1f**). Pre-Art and Art express *Klf2* and *Klf4*, genes induced by laminar shear flow and preferentially associated with linear segments compared to branched vessels^13,14^.

Venous EC, comprising HEC and non-HEC (Vn) subsets, share expression of the vein- specifying transcription factor *Nr2f2* (Coup-TFII)^15^, and the vein-associated chemokine interceptor *Ackr1* (*Darc*^16^; **Fig. 1f**). HEC express genes required for lymphocyte recruitment including *Chst4* and *Glycam1*^17,18^, with more pronounced expression on distal HEC in the venous branch (late HEC; **Fig. 1e-f**). Consistent with their large size and plump morphology and with prior whole genome expression studies of sorted HEC^6^, their gene signature is enriched for glycoprotein synthesis (not shown), and they have uniquely high numbers of transcripts per cell (**Fig. 1g**).

Cells of the Vn subset branch prominently from proximal HEC and TrEC in tSpace projection (**Fig. 1e**). To identify these endothelial cells in the LN vasculature, we imaged whole LN removed shortly after i.v. delivery of fluorescently labeled antibodies (Methods): anti-Ly6c, specific for arteries and capillaries; antibody MECA79 to the Peripheral Node vascular Addressin (PNAd) defining HEV; and anti-PLVAP, which stains capillary and venous EC but not arteries (**Fig. 2a**). Subset markers allow visualization of arterial entry from the LN hilus, linking to capillary arbors in the cortex which in turn connect to HEC. We identified PNAd^-^ veins downstream of and as a continuation of HEV in the lymph node medulla (**Fig. 2a**). PNAd^-^ medullary veins bound injected antibodies to VE-cadherin, PLVAP, and ICAM1 (**Fig. 2b**) but were negative for capillary markers Ly6c and podocalyxin (PODXL; **Supplementary Fig. 4)**, as predicted by gene expression (**Fig. 2c**).

**Figure 2.**
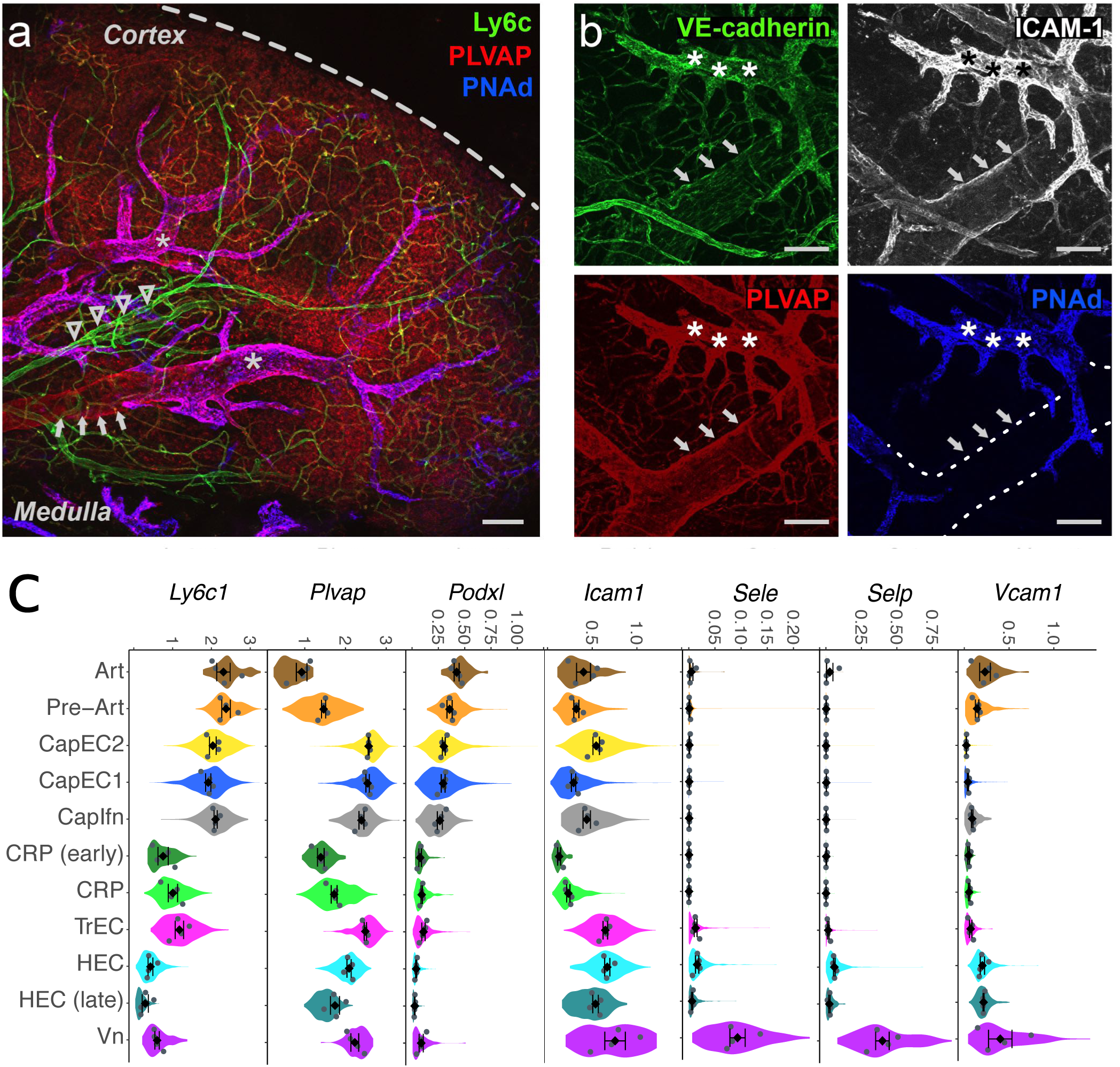
Marker gene expression and immunolocalization of the major arterial and venous populations. Immunofluorescent visualization of PLN vessels using intravenously (i.v.) injected antibodies: anti-Ly6c (green), anti-PLVAP (red), and anti-PNAd (blue). Dashed line, lymph node capsule (a); anti-VE-cadherin (green), anti-ICAM-1 (white), anti-PLVAP (red), and anti-PNAd (blue) (b). Arrow heads, artery (Podxl+ Ly6c+ PNAd-, PLVAP-, ICAM1low). Arrows, medullary veins (PNAd-, ICAM1+, PLVAP+, Ly6c-, Podxl-). Medullary veins are downstream of HEC. Asterisks, HEV (PNAd+). Bars, 100 μm. Dotted line, medullary vein. (c) Violin plots showing expression of genes Ly6c1, Plvap, Podxl and Icam1 corresponding to immuno-stained marker proteins; and Sele, Selp and Vcam1 illustrating selective expression by non-HEV vein. Note the decline in Podxl expression from artery to pre-Art to capillary EC subsets, and a corresponding decline in intensity of staining for PODXL as arteries bifurcate into capillaries in situ in (a). Mean expression values for each of the four independent cohorts (grey dots) and mean and SEM of the cohort means are also plotted (black diamonds) within the violin plots.

The Vn gene signature includes genes associated with regulation of neutrophil activation (GO:1902563; **Fig. 3a**) and platelet degranulation (GO:0002576; **Supplementary Fig. 3**). Vn express Von Willebrand Factor (*Vwf*), which is stored in Weibel-Palade bodies and is released during inflammation to promote platelet adhesion and hemostasis ^19^. Surprisingly, Vn EC lack HEC-associated genes for naïve lymphocyte recruitment (*Chst4, Fut7, Ccl21)*, instead expressing genes for vascular E- and P- selectins (*Sele* and *Selp*) and adhesion receptors *Icam1* and *Vcam1* (**Fig. 2c**) which mediate myeloid cell recruitment. Neutrophils and monocytes are normally excluded from LN homing, but they enter LN’s in large numbers in response to acute inflammation and play an important role in preventing pathogen spread. Prior studies have characterized inflammatory changes in HEC which enable recruitment of myeloid cells along with lymphocytes^20–22^; but the role of medullary Vn has not been examined. We induced inflammation by footpad injection of the bacterial pathogen *S. Aureus* in mice with green fluorescent protein (GFP) expressing neutrophils (LysM^GFP^ mice). One hour later, mice were transfused with CMTPX-labeled lymphocytes and recruitment was quantified by live two photon imaging of the popliteal LN. PNAd^-^ medullary venules exhibited massive and exclusive recruitment of GFP^+^ myeloid cells, contrasting with both lymphocyte and induced myeloid cell interactions in the HEV (**Fig. 3b-e**). In contrast to recruitment through HEV which is PNAd-dependent^21^ myeloid recruitment was robust even in the presence of blocking concentrations of anti-PNAd (not shown), Anti-P-selectin significantly inhibited GFP^+^ cell accumulation on Vn, and combined inhibition of the vascular E- and P-selectins largely abrogated medullary vein interactions of myeloid cells (**Fig. 3f**), even though HEV recruitment was unaffected^21^. We conclude that venules in the medullary environment have a unique phenotype and function, selectively recruiting myeloid cells to the medulla in acute inflammation. The results illustrate a surprisingly localized endothelial cell programming for differential leukocyte recruitment.

**Figure 3.**
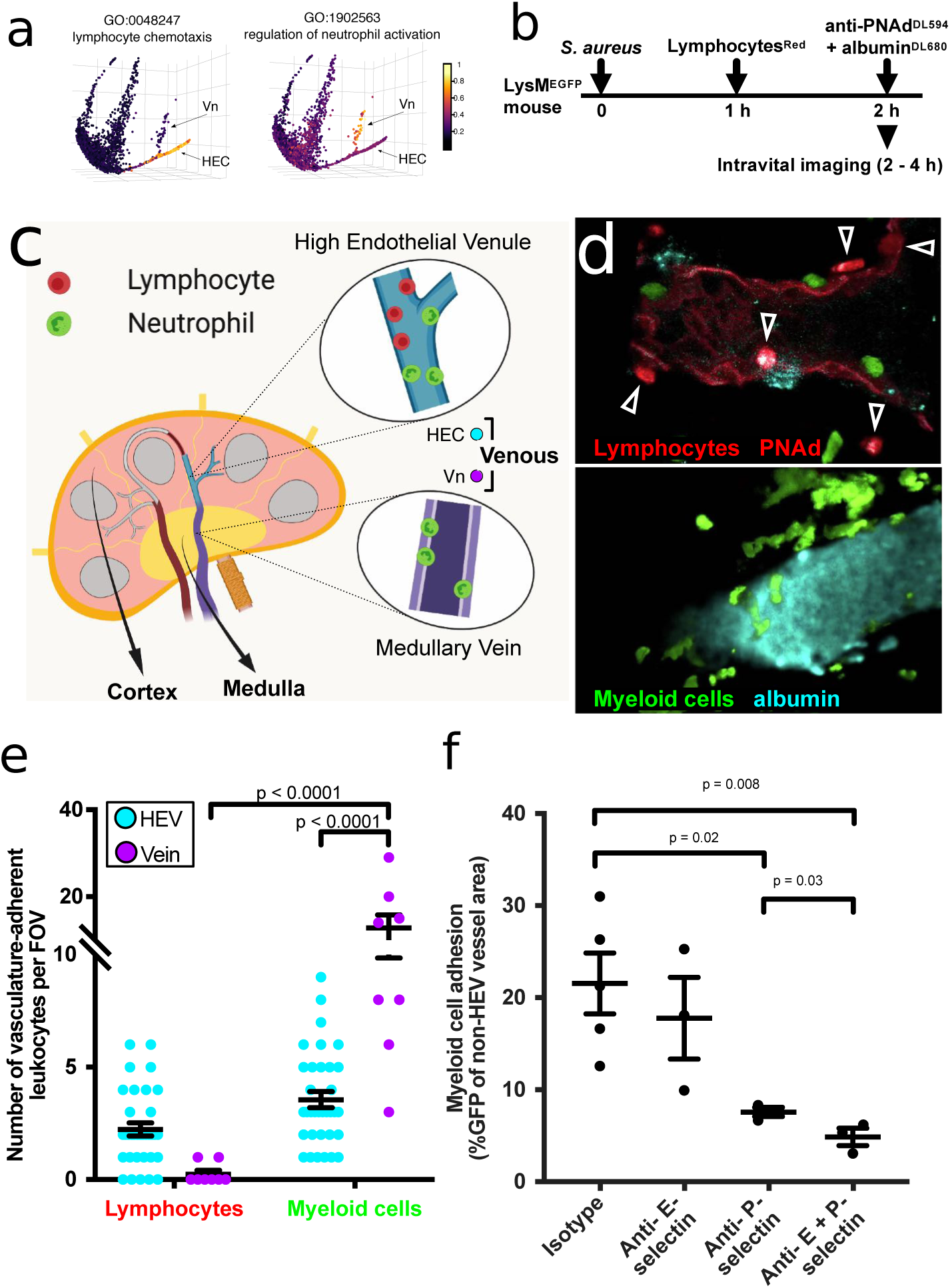
Medullary veins recruit myeloid cells but not lymphocytes in acute inflammation. In situ lymphocyte and myeloid cell recruitment in HEV versus medullary veins of S. aureus infected LysMGFP mice. (a) Pooled expression values of genes from the indicated GO terms (color scale) plotted along the tSpace projection from Fig 1e. (b) Experimental protocol for S. Aureus injection and visualization of lymphocyte and myeloid cell trafficking in LN. LysMGFP recipients received 2.5 x107 S. aureus in the footpad. 1 hour later mice were injected i.v. with CMTPX-labeled lymphocytes (red). The draining LN was imaged from 2 hours to 4 hours post infection using two-photon videomicroscopy. (c) Schematic depicting the location of HEV and medullary veins visualized. (d) Representative fields of view from 2 photon videomicroscopy of a LN from mice treated according to (b). Myeloid cells (green) and lymphocytes (red; arrow heads) arrested in HEV (upper panel) or medullary vein (lower panel). HEV, identified by injection of red fluorescent anti-PNAd at a non-blocking concentration immediately prior to sacrifice, are readily distinguished from migrating lymphocytes and from PNAd- medullary veins. Venular lumen is highlighted by Dylight-680 labeled albumin (cyan). (e) Quantification of lymphocyte and myeloid cells adherent to HEV and medullary veins. n = 34 fields of view (FOV) for HEV and 8 FOV for veins from a total of 4 mice. Data shown as mean ± SEM. (f) Inhibition of myeloid cell accumulation in medullary veins by antibody blockade of P- and E- selectin. Test or isotype control antibodies were injected i.v. 20 min before footpad S. aureus infection in LysMGFP mice, and draining LN visualized 2 hours post infection. Myeloid cell (GFP+) adhesion to medullary veins was quantified from 24 - 47 FOV of popliteal LN over 2 hours of imaging. Each point represents an average of the values collected from one mouse. n = 5 mice for isotype, n = 3 mice for all other groups. Data shown as mean ± SEM.

### Characterization of capillary subsets

Five capillary populations shared expression of *Cdh13*, *Emcn*, *Gja1* and *Gpihbp1,* previously described as gene markers of capillary EC in PLN^6^ (**Fig. 1f**). One capillary cluster comprises transitional phenotype capillary EC (TrEC). TrEC express canonical capillary genes as well as some HEC genes, albeit at low levels compared to *bona fide* HEC (**Fig. 4a, c**). They bridge other CapEC to the venous branch in tSpace projections (**Fig. 1e**), further suggesting a close relation to HEC. TrEC express *Chst2* and the HEC genes *St3Gal6* and *Fut7* which encode glycosyltransferases for the synthesis of sialyl LewisX (SLex) and 6-sulfo-SLex, carbohydrates that can initiate tethering of lymphocytes under shear flow^18^. Synthesis of PNAd, the mature multivalent L-selectin ligand for lymphocyte homing that defines HEV also involves *Chst4* and requires the core 2-branching enzyme encoded by *Gcnt1. Chst4* and *Gcnt1* are nearly undetectable in TrECs, suggesting that TrEC and HEC might display different glycotopes. Thus, we used antibodies to SLex and PNAd to identify TrEC *in situ.* Imaging revealed a significant population of BEC that co-stained for SLex and for capillary antigens (**Fig. 4b, Supplementary Fig. 5**) but lacked mature PNAd. They were morphologically thin-walled and were found immediately upstream of HEV, correlating with their position in trajectory space (**Fig. 1e**).

**Figure 4.**
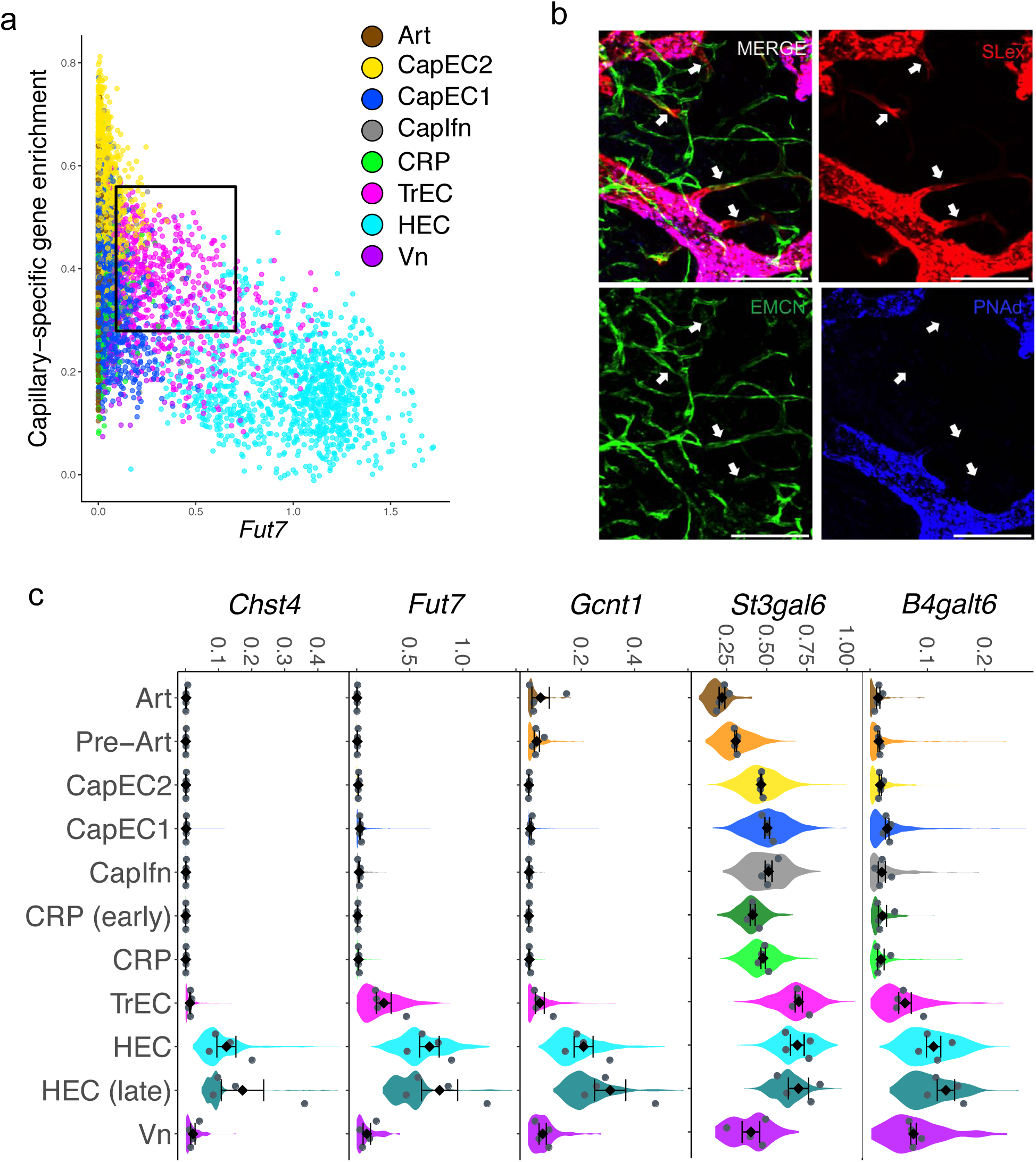
Transitional phenotype capillary EC occupy capillary-HEC junctions. (a) Scatter plot of cells showing Fut7 expression by capillary EC defined by an enrichment score for capillary-specific genes. Cells colored by major cell type. (b) Immunofluorescence image of PLN with intravenously injected anti-SLex (red), anti-PNAd (blue), and anti-capillary (EMCN; green) antibodies. Scale bar 100 µm. Arrows point to Slex+ EMCN+ PNAd- TrEC. (c) Expression of Chst2, Fut7, Gcnt1, St3gal6, B4galt6 in the BEC subsets. Violins show the expression distribution of all cells. Mean expression values for each of the four independent cohorts (grey dots) and mean and SEM of the cohort means are also plotted (black diamonds).

CapEC1 and CapEC2, the two most abundant capillary populations (**Fig.1c**), are centrally located among EC in trajectory space, acting as a hub from which differentiated venous and arterial branches emerge (**Fig. 1e**). The CapEC1 gene signature is enriched for Type IV collagen trimers (GO:0005587; *Col4a1*, *Col4a2*) as well as *Igfbp3* involved in maintaining vascular tone and blood pressure^23^. They have high vascular endothelial growth factor receptor activity (GO:0005021), yet also express negative regulators of angiogenesis (*Hlx* and *Igfbp3*) and have enhanced expression of genes *Id1* and *Id3* encoding inhibitor of differentiation proteins, bHLH transcriptional regulators that restrain cell differentiation and delay cellular senescence^24,25^. They are enriched for genes for negative regulation of leukocyte tethering (GO:1903237)^26,27^ (**Fig. 1f, Supplementary Fig. 3**) .

CapEC2 express *Egr1, Cxcl1* and genes reflecting NFkB activation (*Nfkbia, Nfkbid, Nfkbiz*) and JNK activation (*Jun, Junb, Jund, Fos, Fosb, Atf3*). These genes and pathways are induced by spatial gradients in fluid shear stress^28–31^. ^32,33^. Compared to CapEC1, CapEC2 also show reduced Gja4 and Id1 and enrichment for genes in the MAPK cascade (GO:0000165) and WNT signaling (GO:0060070; **Supplementary Fig. 3**), characteristics that parallel findings from a ChIP-seq-based analysis of a disturbed oscillatory shear stress response^31^. These CapEC2 features suggest exposure to gradient or disturbed shear stress.

A third relatively rare population, CapIfn, has a prominent signature of interferon signaling (GO:0060337), with high expression of transcription factor *Irf7*, interferon response genes *Ifit1*, *Ifit2*, and chemokines *CXCL9* and *10 (***Fig. 1f**). CapIfn also express *Isg15* and *Gadd45a,* genes associated with an “apoptotic-like” signature described in other single cell studies^34^.

### An angiogenic capillary subset enriched for stem cell-associated genes

Unsupervised clustering identified a distinctive population of activated capillary endothelial cells, CRP, that occupy their own branch in trajectory space, linking to CapEC1, 2 and TrEC (**Fig. 1e**). Their gene expression signatures reveal a progenitor-like phenotype. They have uniquely high expression of *Ets2* and *Sox7*, encoding transcription factors implicated in early specification of endothelial cells during development^35,36^ and they display genes and features of stem or progenitor cells in other systems. These developmental EC genes are additionally enriched in early vs late CRP as defined by cell positions along their tSpace branch. CRP also express genes associated with neural and hematopoietic stem or progenitor cells including *Cxcr4*^37^*, Nes*^38^*, Kit*^39^*, Lxn*^40^, and *Sox4*^41^; and they are enriched in spliceosome genes (GO:0097525) including *Snrpa1* and *Snrpd1* (**Fig. 5a**), which participate in the acquisition and maintenance of pluripotency in embryonic stem cells ^42,43^. Like embryonic and neural stem cells, CRP uniquely lack expression of *Neat1*, a long non-coding gene widely expressed in differentiated cells for paraspeckle assembly and double stranded RNA processing during stem cell fate selection ^44,45^ (**Fig. 5a**). Multipotent stem and progenitor cells have more diverse gene and protein signaling than their differentiated progeny, reflecting their developmental plasticity. These characteristics can be quantified by calculating the ‘signaling entropy rate’ using the SCENT algorithm^46^. Early CRP display higher entropy than other EC subsets, while Art and late HEC subsets display the lowest entropy (**Fig. 5b**). Dividing cells among CapEC1 share some of these correlates of potency, including high entropy; however, cells with the highest entropy tend to map to the ‘origin’ of the CRP branch in trajectory space (**Fig. 5c**). Finally, CRP express *Angpt2*, *Apln*, *Esm1*, *Nid2*, *Pdgfb* and *Pgf* (**Fig. 1f**), genes characteristic of angiogenic tip cells, precursors to new vessels formed during sprouting angiogenesis^47,48^. Together these observations suggest that CRPs comprise an activated or primed capillary population with the potential to contribute to vascular maintenance and vessel growth. Consistent with this, CRP are enriched in cells with genes for cell division (**Fig 5d**). Although only ∼10% of CRP have dividing cell gene signatures, in resting LN ∼60% of all cells with gene signatures for cell division align with CRP, with most others distributed to CapEC1 and TrEC (**Fig 5e**). Interestingly, division increased as *Apln* declined in the transition from early to late CRP and to TrEC, which are *Apln-* (see below).

**Figure 5.**
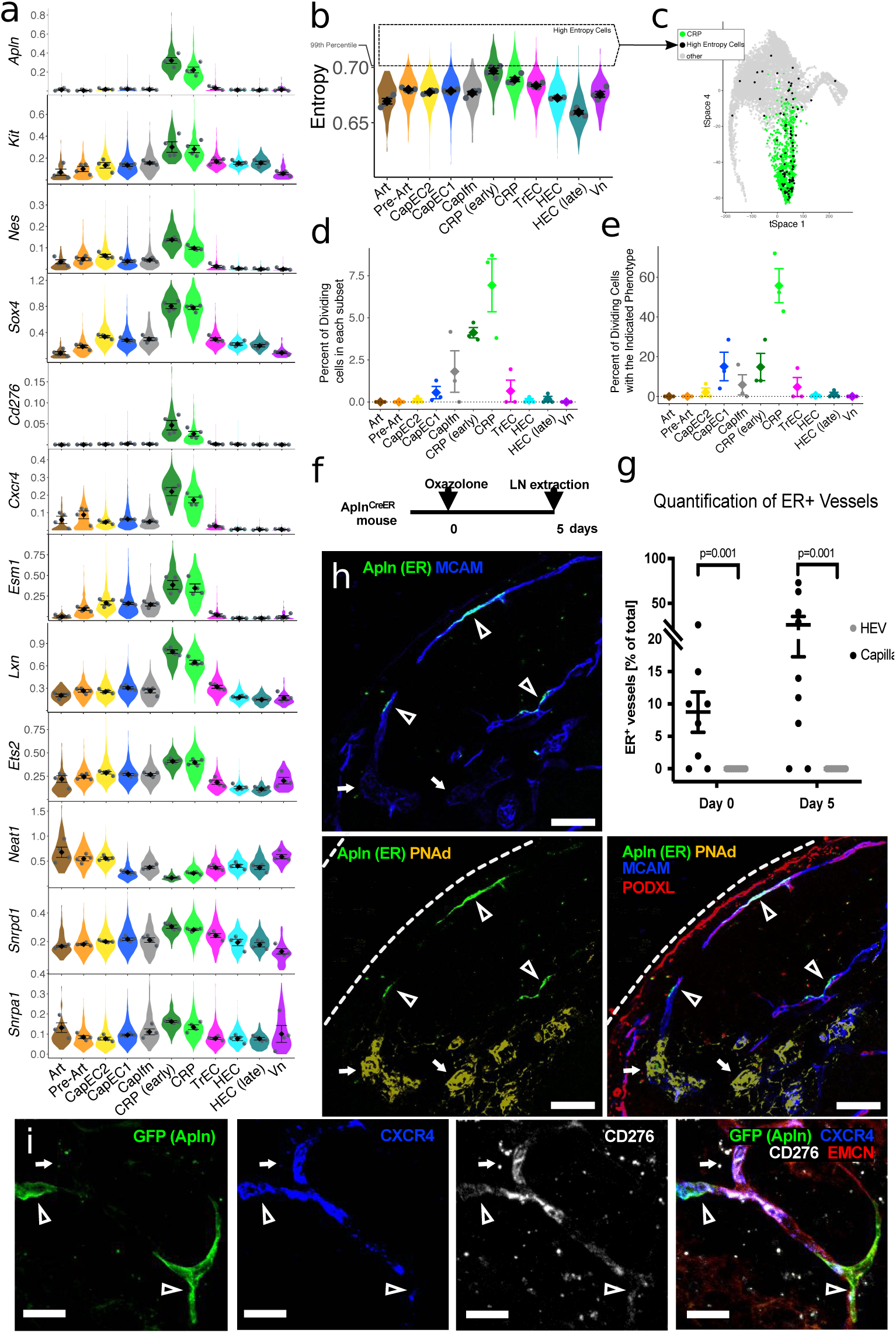
Stem cell features, markers and immunolocalization of a capillary resident regenerative population (CRP). (a) Expression of selected stem or progenitor cell related genes. (b) Signaling entropy rate (entropy) for each BEC subset. Highest 1% of all BEC, dashed box. High entropy cells are enriched in early CRP. (c) tSpace projection from Supplementary Fig. 1 with all cells. CRP, green. High Entropy cells (top 1% as gated in (b)) are black. Other cells, grey. Interactive rendering available: https://stanford.io/2WXR811 (d) Percent of each BEC subset classified as dividing based on high pooled expression of cell cycle genes. Points represent values for individual samples. Diamonds, average of all samples. Error bars, standard error of the mean. (e) Phenotype of dividing cells presented as percent of dividing cells with the indicated BEC phenotypes. (f) Experimental timeline for (g) and (h). (g) Quantification of ER+ endothelium by immunofluorescence histology in resting (day 0) PLN and in PLN (day 5 after) cutaneous inflammation. Expressed as ER+ capillaries (PODXL+ or MCAM+ PNAd- EC) or ER+ HEV (PNAd+) as percent of capillary EC or HEC scanned. Units are lengths of vessel segments in arbitrary units. No ER+ HEC were detected out of over 5000 scanned in resting and over 5000 in inflamed PLN. (h) Representative images of resting PLN from AplnCreER mice stained with anti-PNAd (yellow), anti-PODXL (red), anti-ER (green) and intravenously injected anti-MCAM (blue). Arrow heads point to ER+ CRP. Bars, 50 μm. (i) Representative images of resting PLN (72h post 4-OHT) from AplnCreER, mTmG mice stained with injected anti-CXCR4 (blue), anti-CD276 (white) and anti-EMCN (red). Arrow heads point to GFP+ cells in capillaries. Bars, 20 µm.

As CRP selectively express *Apln* (**Fig. 5a**) we localized CRP within the LN vasculature of Apln^CreER^ mice (**Fig. 5f**) by staining for the human estrogen receptor (ER), which serves as a surrogate of Apln expression in these mice^49^. ER^+^ EC were readily visualized as thin-walled endothelial cells in capillaries. HEC were not stained by anti-ER, and importantly ER expression detectable by immunofluorescence histology remained restricted to capillary EC after immune challenge (**Fig. 5g, Supplementary Fig. 6**). ER+ CapEC were rapidly labeled by intravenously injected antibodies to surface antigens, consistent with luminal contact and integration into the capillary endothelium (**Fig. 5h**). *CD276*^50^and *Cxcr4*^48^, known markers of angiogenic potential in EC, are also selectively expressed by CRP (**Fig. 5a**). Anti-CD276 and CXCR4 antibodies injected intravenously labeled a subset of EC limited to capillaries. In Apln^CreER^ x Rosa26^mTmG^ (Apln^CreER; mTmG^) mice pulsed three days previously with 4-Hydroxytamoxifen (4-OHT) to induce GFP reporter expression in AplnER+ cells, GFP^+^ CapEC showed heterogenous staining for injected anti-CD276 and CXCR4 (**Fig. 5i**).

To assess their fate, we immunized Apln-CreER x R26 mTmG mice with Complete Freund’s Adjuvant (CFA) 24 hours after injection of the short acting tamoxifen metabolite 4- hydroxytamoxifen (4-OHT; serum half-life 6 hours^51^). Three and a half weeks later, many HEC and capillary EC were positive for the reporter, confirming EC neogenesis from AplnERTCre- expressing precursors (**Fig. 6).** Similar results were obtained in a repeat experiment in which 4OHT was administered 3 days prior to immunization (**Supplementary Fig. 7**) of mice in one leg: GFP expression remained concentrated in scattered CapEC in the control LN even 3 weeks later, whereas numerous GFP^+^ progeny incorporated into HEV in the immunized node. Finally, consistent with the restricted detection of *Apln-*promoter*-*driven ERTCre in CapEC even after immunization (**Fig. 5f-g**), we found that reporter was selectively induced in capillary EC even when 4-OHT was administered 24 hours <u>after</u> immunization: Incorporation of reporter+ progeny was seen in HEV examined 12 days later (**Supplementary Fig. 8**). In conjunction with their expression of genes involved in embryonic vascular development and their enrichment in cells undergoing basal cell division, the results suggest that CRP are a poised regenerative subset that can contribute to vessel maintenance at steady state and to neogenesis of EC including HEC during LN angiogenesis.

**Figure 6.**
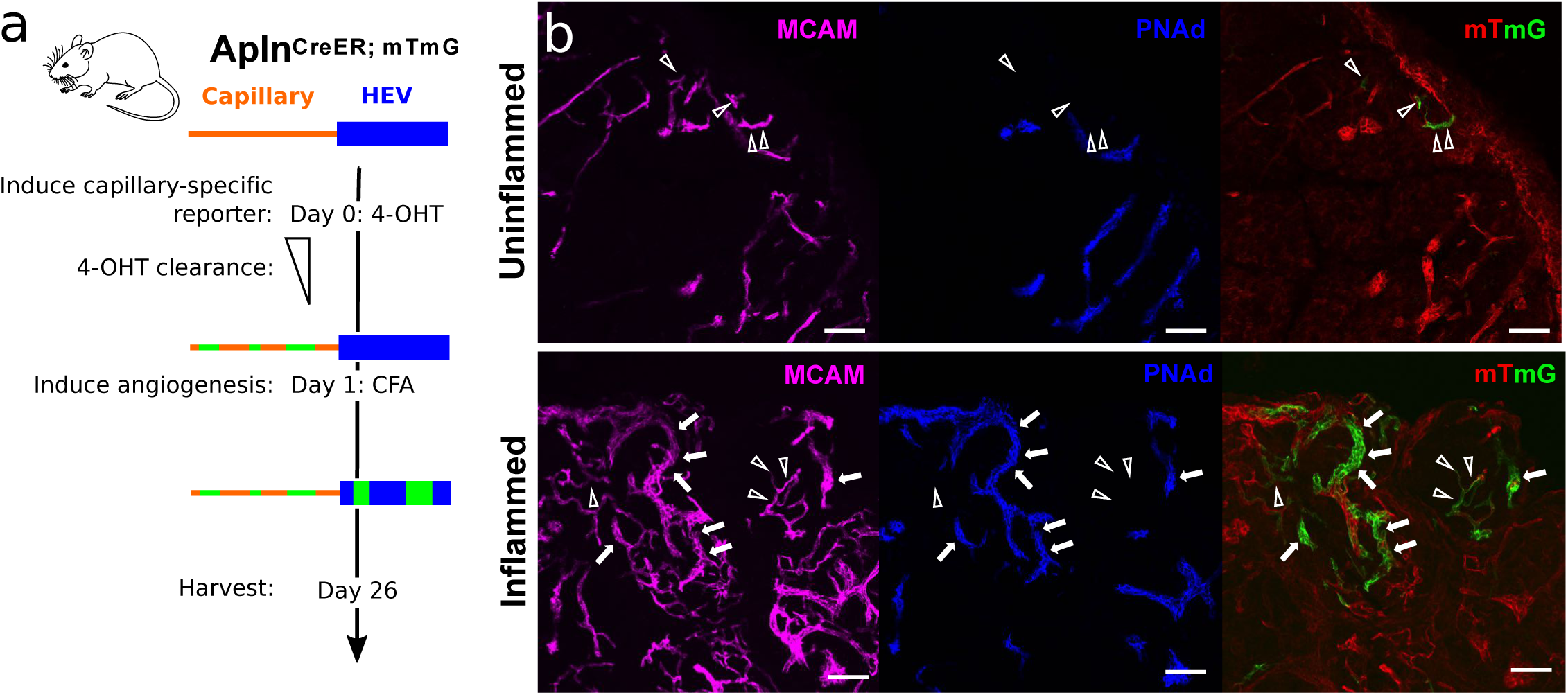
Lineage tracing of AplnERTCre-expressing capillary EC. (a) Experimental protocol. (b) Representative images of Apln-driven reporter expression (GFP) in PLN from Apln-CreER x R26-mTmG mice at rest (upper) and three and half weeks post- immunization with CFA (lower panel). EC subsets were labeled by i.v. injection of the indicated antibodies 10-20 minutes before sacrifice: PNAd (blue), MCAM (magenta). Arrow heads, capillaries. Arrows, HEV. Bars, 50 µm.

We recently identified an *Apln*-expressing metaphyseal EC subset in the adult bone marrow that contributes to the hematopoietic stem cell niche^52^: fate mapping in Apln-CreER x Rosa26- mTmG mice suggested that the Apln+ EC contribute to endothelial homeostasis and neogenesis in the marrow vasculature, as shown here for LN CRP. We therefore sought to identify CRP-like EC in BM and other tissues, taking advantage of scRNAseq profiles of BEC from the Tabula Muris (TM) consortium^53^ and the gene expression omnibus (GEO)^54,55^. We used mutual nearest neighbor analysis to perform a global alignment of ∼38,000 BEC, including cells of our PLN2 and 3 samples, to cells of our reference PLN1 sample (Supplementary Figure 9a). We defined candidate CRP-like cells as cells whose gene expression profiles 1) correlated more highly with the mean gene profile of CRP than with other CapEC or differentiated subsets; and 2) aligned with PLN CRP in UMAP projection. A small percentage (0.3-13%) of BEC met these criteria in most tissues (Supplementary Figure 9b): the majority of these expressed *Apln* as well as other CRP genes including *Cd276, Cxcr4, Lxn, Mcam* and *Trp53i11* (Supplementary Figure 9c; Supplementary Table 2). The CRP-like EC express many angiogenic tip-cell related genes, and like LN CRP share high entropy compared to differentiated BEC in their respective tissues (Supplementary Fig 9 c-d). In addition to aligning with LN capillary CRP, most CRP-like cells express capillary-associated gene markers including *Gphibp1, Ly6c1* and *Cdh13*, and lack venule and artery genes (not shown). However, BM CRP-like cells had lower *Ly6c1* than cells in other sites. CRP-like EC were extremely rare or absent in liver. In lung, *Apln* is highly expressed by capillary aerocytes, but these cells, which are unique to the pulmonary vasculature, are otherwise unrelated to CRP. The results suggest that CRP are a rare but widely distributed capillary phenotype angiogenic population.

### Gene regulation along cellular trajectories

We next examined changes in genes and cell features along cellular trajectories. We isolated cells along paths from early CRP to arteries or to HEC, or from CapEC along the venous branch leading to Vn (**Fig. 7a**), and visualized expression of genes or gene modules by cells along the trajectories (**Fig. 7b-e, Supplementary Fig. 10**). The heatmap illustrates enrichment of cells with high cell cycle scores between or along the path from early CRP to CapEC and TrEC, peaking in correspondence with late CRP or the transition to TrEC (**Fig. 7b, Supplementary Fig. 10a**). Conversely *Apln* declines rapidly in the transition from early to late CRP and TrEC, and consistent with this dividing CRP have reduced *Apln* expression compared to early CRP (**Fig. 7b, Supplementary Fig. 10a**). CapEC on the trajectory to Art express genes implicated in developmental arteriogenesis^56^. These include *Sox17*, *Nrp1, Gata2, Klf2,* and genes for Bone Morphogenic Proteins (BMP), Notch and Ephrin signaling components and downstream targets (*Acvrl1, Tmem100, Msx1, Notch4 and Notch1, Jag1, Jag2, Hey1, Efnb2*) that program the mature arterial phenotype (**Fig. 7e**). Features reflecting laminar shear stress (*Klf2, Klf4*, GO:0034616; **Fig 7e and f**) increase along the trajectory to Art, while the oscillatory or gradient shear stress signature peaks in capillary EC, especially the CapEC2-rich region of the trajectory leading to arterial EC (*Cxcl1*, *Egr1*, OSS; **Fig. 7c, e and f**). Genes involved in gas and metabolite exchange *(Aqp1*^57^*, Car2 and Car4)* are high in CapEC and lost in the progression to Art. Surprisingly, arterial EC express key genes for elastin fiber assembly (*Fbln5, Eln, Ltbp4 and Lox*), suggesting that arterial EC may directly participate in the assembly of the inner elastic lamina between EC and smooth muscle cells (**Fig. 7e**). The CRP to Art trajectory also displays genes involved in arterial EC progenitor migration in development, discussed below.

**Figure 7.**
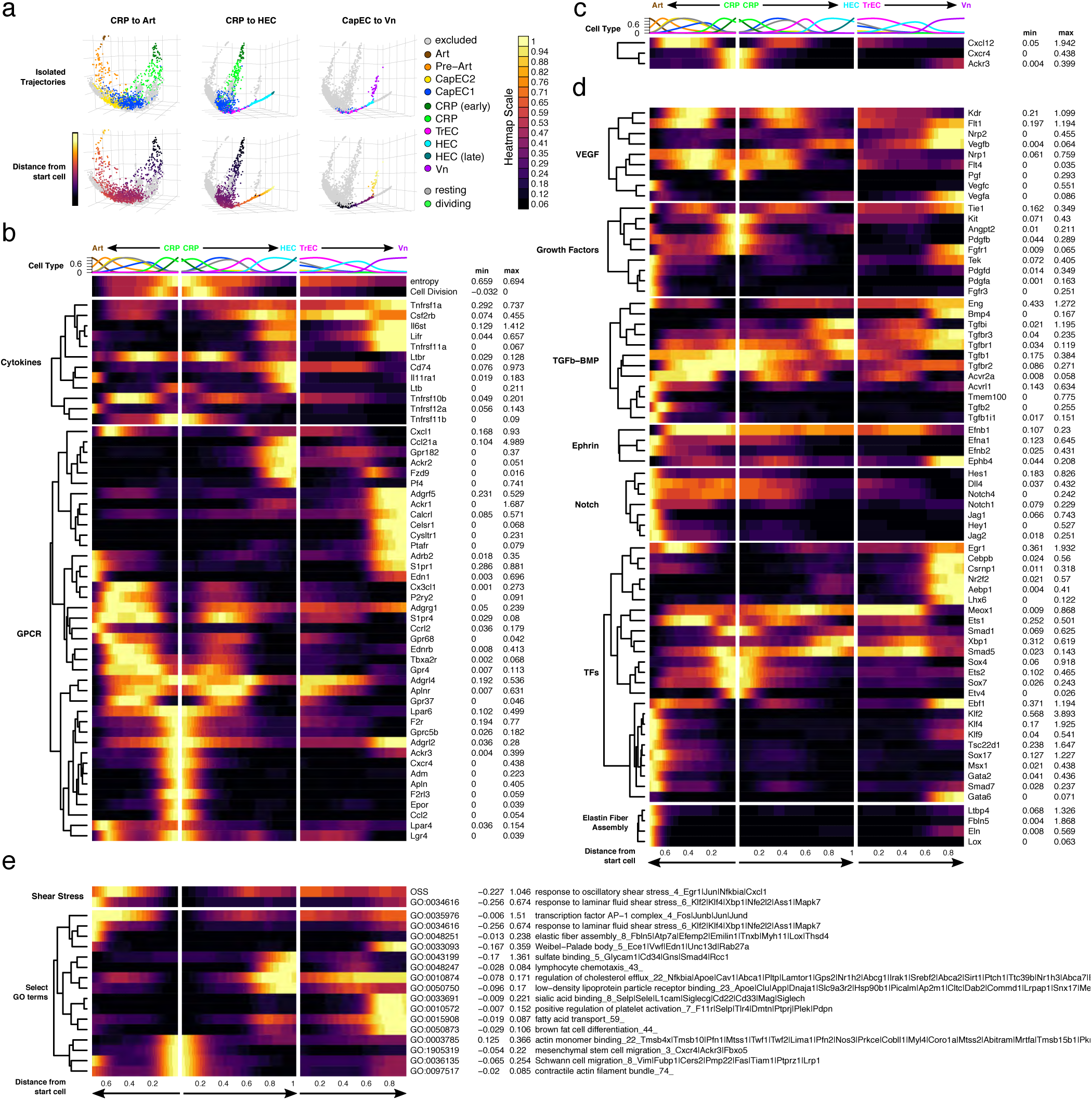
Trajectories align EC subsets and mechanisms of EC development and specification to the vasculature. (a) Cells along KNN-based trajectories were isolated (Methods). (b - e) Expression of selected genes and gene set enrichment scores along cell trajectories from mature CRP to Art (plotted leftward), and from CRP to HEC or Vn (rightward). Cells along the trajectories were manually gated in the first 5 principal components of trajectory space and aligned according to distance from early CRP. Representation of cell types along the trajectories is indicated at top. Normalized count data plotted as a function of trajectory distance was smoothed using a gaussian kernel. Trajectory distances were scaled to the longest trajectory (CRP to HEC). Genes were grouped according to biological class or function. Imputed gene expression values for all cells (without duplications) were calculated independently and used for hierarchical clustering within each gene group. Cxcr4, Cxcl12 and Ackr3 are shown as a separate group at the top in (c) (see results). Average normalized expression values for the min and max subset are indicated to the right of each heatmap. OSS: pooled expression of oscillatory shear stress genes: Egr1, Nfkbia, Junb, Cxcl1. Cell Division: pooled expression of cell cycle genes.83

In contrast, cells along the trajectory from CRP to Vn or HEC express determinants of venous fate and phenotypes. *Nr2f2* encodes CoupTF2, a TF required for venous differentiation in development: *Nr2f2* is first expressed in TrEC and is maintained along the venous trajectory. Notch is inhibited during venous differentiation^15^, and Notch signaling components are downregulated early along the venous trajectory (**Fig. 7e**). The lymphotoxin receptor LTBR is required for HEV development and maintenance. *Lbtr* is broadly expressed, but lymphotoxin beta (*Ltb*), a component of HEV-specifying lymphotoxin dimers, appears in the late HEC portion of the venous axis, suggesting the potential for autocrine signaling to induce and maintain the HEC phenotype (**Fig. 7c**). In addition to mapping known mediators and mechanisms of arterial and venular specification to cellular trajectories, the analysis identifies novel candidate transcription factor genes that may contribute to specialization of large vessels (e.g. *Ebf1*, *Klf9*), arteries (e.g. *Tsc22d1*), HEC (e.g. *Xbp1*, *Meox1*), HEC and vein (*Aebp1*) or veins (*Lhx6*, *Csrnp1*, *Gata6*) (**Fig. 7e**). For example, *Gata6* enhances Tumor Necrosis Factor-alpha induced VCAM1 expression in EC cultures^58^ and its expression by TrEC and Vn may thus contribute to selective Vn expression of *Vcam1*.

G protein linked receptors (GPCR) serve as environmental sensors. With their ligands GPCRs regulate vascular development, endothelial function and migration. We examined expression of GPCR along the aligned trajectories. The vasodilatory flow sensor gene *Gpr68*^59^ first arises in pre-Art capillary EC and is retained in Art (**Fig. 7c**). *Cysltr1*, encoding a sensor for myeloid cell-derived leukotrienes^60^, is upregulated in the transition to Vn (**Fig. 7c**). Genes encoding thrombin and erythropoietin receptors (*F2r* and *Epor*), sensors that drive angiogenesis and vasculogenesis^61,62^, are expressed selectively by CRP, suggesting sensitivity to local thrombotic or inflammatory protease activity and oxygen deficiency (**Fig. 7c**). Patterns of expression of genes for *Cxcl12* (in CapEC and pre-Art), and for its chemoattract receptor *Cxcr4* (expressed by CRP and subsets of CapEC1 and 2 along the trajectory to Art) suggest retention of a developmental programs for tip cell migration along capillaries into developing arteries (**Fig. 7d**), as observed during retinal arteriogenesis^63^. The CXCL12 molecular sink receptor ACKR3, whose expression in trailing cells establishes a CXCL12 gradient for directed germ cell migration in zebrafish^64^. *Cxcr4* is expressed by CRP and some CapEC in the arterial branch suggesting a parallel role. These observations suggest that CRP may be primed for migration. Consistent with this, CRP are enriched in genes for cell locomotion, extracellular matrix remodeling, actin assembly and disassembly and lamellipodium formation (**Fig. 7c**); but they also express genes for known inhibitors of migration and sprouting behavior including *Arhgap18*^65^ and *Csnk2b*^66^ (**Supplementary Table 1**).

## Discussion

The vascular endothelium plays a central role in lymphoid tissue development and function. Our survey of the LN blood vascular endothelium defines the diversity of EC phenotypes and identifies novel cell subsets and functions. We uncover multiple subsets of capillary endothelial cells with distinct gene expression including TrEC, a transitional phenotype subset intermediate in gene expression and in physical location between other CapEC and HEC. We show that this subset expresses glycotopes^18^ for lymphocyte tethering under flow. Presentation of tethering glycotopes by capillary EC immediately upstream of HEC may facilitate lymphocyte homing by allowing lymphocytes to initiate interactions with the endothelium prior to entering HEV. We identify a unique profile of medullary venous EC and show that they explicitly recruit myeloid cells and not lymphocytes to the LN medulla in response to acute bacterial challenge. Neutrophils block pathogen spread beyond the initial tissue draining LN^21^, and neutrophils recruited into the medulla may be well positioned to intercept bacteria prior to their exit into the efferent lymphatics. Finally, we identify a primed capillary resident population, CRP, that display features associated with multipotent progenitor cells and participate in basal endothelial proliferation and in vascular neogenesis in response to immunization.

New endothelium in physiologic angiogenesis is thought to arise principally from local EC^67^, and circulating endothelial progenitors do not contribute to new endothelium in immunized LN^4^. Prior studies have highlighted multipotent vessel-resident EC progenitor populations in large vessels^68,69^ and lung^70^ and precursors in developing bone^71^ that can differentiate into diverse EC phenotypes. CRP appear distinct from these populations: They lack *Bst1* and *Procr*, markers of resident endothelial progenitors described in large vessels of liver^68^ and fat pad^69^. They express *Kit*, a marker of clonally proliferative EC progenitors in lung^72^; but published single cell datasets of lung blood vascular EC include only extremely rare EC with CRP-like gene profiles. CRP are also distinct from angiogenic tip cells in that CRP are in direct contact with the lumen and are integral to the vessel lining, whereas tip cells lack luminal contact, instead leading the blind end of invasive sprouts^73^. However, CRP share gene expression and precursor potential with tip cells, and thus may be primed for tip cell behavior or alternatively for intussusceptive (splitting) angiogenesis. CRP appear closely related to *Apln*-expressing EC we have observed in the adult bone marrow, which fate mapping suggests contribute to irradiation-induced neogenesis of endothelial cells including arterial EC in the marrow compartment^52^. Moreover, we show that computational alignments and shared gene signatures identify rare CRP-like capillary phenotype EC in many tissues. Thus CRP may contribute widely to vascular homeostasis, providing a distributed pool of regenerative cells for local vascular maintenance and replenishment. Consistent with this thesis, a recent scRNAseq study also reported rare *Apln*+ “angiogenic” EC in normal tissues^74^.

Interestingly, in PLN and in most tissues profiled *Apln* expression appears quite selective for CRP or CRP-like EC. At the gene level, its expression is highest in ‘early’ CRP which also show the highest signaling entropy among the BEC profiled here. *Apln* is downregulated progressively in cells along the trajectory from early to late CRP and CapEC; and its translation to protein as assessed by *Apln promoter-*driven expression of Cre-Ert2 remains restricted to capillaries even in immunized lymph nodes, in which many HEV and other BEC are undergoing active proliferation. Thus Apelin is not a general marker of EC activation or proliferation. These considerations emphasize the similarities between early CRP and tip cells in models of sprouting angiogenesis: tip cells like early CRP are *Apln* high, predominantly non-dividing and display signatures of cell migration; whereas cell division occurs primarily among trailing ‘stalk’ cells^75^.

The migratory signature of CRP suggests that they may have the capacity to crawl within capillaries to contribute to new vessel formation. We show that CRP express genes for locomotion including a *Cxcr4* chemoattractant program with the potential to drive precursor migration toward pre-artery-expressed Cxcl12 for arteriogenesis, as observed for tip cell progeny in retinal development^76^. We also show CRP expression of *Ackr3* encoding the Cxcl12 interceptor Ackr3 (Cxcr7): Ackr3 internalizes Cxcl12, and in developing cell systems can enhance Cxcr4-driven directional migration by reducing chemoattractant levels at the “rear” of a migrating population^77^.

CRP display sensory receptors for angiogenesis signals, including thrombin, erythropoietin and VEGFs, and thus appear well programmed to respond to requirements for endothelial proliferation. Consistent with this, we show that nearly half of all BEC with basal cell division signatures align with CRP, predominantly late CRP as just mentioned. CRP may represent a temporary state of activated capillary EC, as suggested by enrichment in dividing cells and similarities to tip cells in their gene expression. Alternatively, they may represent a resident progenitor pool, analogous to regenerative stem or progenitor cells in other settings (i.e. intestinal epithelium and hematopoietic systems). They embody high entropy and express genes characteristic of multipotent progenitors and stem cells. Cell transfer or clonal cell culture studies may help distinguish these possibilities. Our data do not address the precursor potential, in terms of contribution to new vessel formation, of the other EC subsets defined here. CapEC1, TrEC, and early HEC display higher signaling entropy than the most differentiated arterial and venous EC in our samples, suggesting that these EC may also retain developmental plasticity. Indeed, HEC have been reported to contribute to neo-synthesis of HEC and capillary cells when injected into a recipient mouse LN^4^. In other cell systems (e.g. the intestinal epithelium), when subjected to the challenge of injury or stem cell depletion, recently differentiated cells can even re-acquire multipotency and renew stem cell pools^78^: Some subsets of BEC may be capable of doing so as well. The relative contributions of different BEC subsets to neo-synthesis of specialized EC may be a function of the tissues they reside in and the challenges in their micro-environment.

We also observed differences among the major CapEC pool, defining two related clusters, CapEC1 and CapEC2. Both share canonical CapEC genes and genes for gas and metabolite transport which are downregulated in pre-artery and terminal arterial EC subsets. CapEC1 feature high expression of inhibitor of DNA binding proteins (Id1, Id3), proteins that act as inhibitors of cell differentiation. CapEC2 express genes (e.g. *Egr1, Cxcl1*) and pathways (NFkB, Jnk, Wnt, MAPK cascade) induced by oscillatory or gradient shear stress which may occur at vessel bifurcations. Interestingly, these genes decline along a trajectory from CapEC2 to arterial EC, while laminar flow associated genes Klf2 and Klf4 progressively increase. Pre-Art may thus correspond to the arterioles and potentially to cells lining arteriovenous communications in the LN.

The ability of medullary veins to selectively recruit myeloid cells, and the selective expression and role of vascular selectins in the process, reveal strikingly local vascular specialization. Previous studies have shown the inflammatory stimuli can induce *de novo* monocyte and neutrophil recruitment to LN but have focused on the role of HEV and the HEV L-selectin ligand PNAd which recruit cells preferentially into the deep cortex (T cell zones). We showed recently for example that neutrophils home via HEV into *S. aureus* challenged LN in a PNAd-dependent, vascular selectin-independent process^21^. In contrast, we find here that medullary veins lack the machinery for naïve lymphocyte homing, instead selectively recruiting myeloid cells using the vascular selectins. The medulla of the lymph node contains pathogen-trapping lymphatic EC networks and that we have recently characterized at the single cell level^79^: recruitment of neutrophils to the medulla in acute bacterial challenge may contribute to the important role of myeloid cells in reducing pathogen transit into efferent lymphatics and systemic spread of infection.

Our results show that the major EC subsets, defined by gene signatures, map to specific locations within the steady state vasculature. Indeed, alignment of cells along nearest neighbor trajectories appears to recapitulate the overall architectural arrangement of EC in the blood vasculature. Imaging confirms the computationally predicted positioning of TrEC between CapEC and HEV; of CRP within capillary segments; and the branching of PNAd negative veins from HEV. Trajectory analysis also reveals that capillary EC aligned along trajectories to mature arterial EC express transcriptional programs that, in development, support arteriogenesis. Similarly, genes that program developmental specification of veins are retained along the venous branch. This ‘retention’ of artery and venous specifying programs may reflect the continuous steady state replenishment and developmental programming of these specialized subsets from dividing CRP or other precursors. Alternatively, retention of developmental genes could serve to pre-program segmental EC differentiation during the rapid vascular expansion of immune angiogenesis.

As shown here, single cell analysis has the potential to identify EC subsets; elucidate developmental processes, transcriptional and regulatory pathways that program their specialization; and map transcriptional phenotypes to the vasculature, providing a molecular blueprint of the vascular endothelium. Targeting specific subsets and processes defined here holds promise to treat a variety of vascular, immune and inflammatory disorders through manipulation of angiogenesis and immune responses.

## Acknowledgments

We thank Nicole Lazarus for technical advice, Dhananjay Wagh for single cell sequencing, and Karen Hirschi for critical review. This work was supported by NIH grants R01 AI130471 and R37 AI047822 and award I01 BX-002919 from the Dept of Veterans Affairs to ECB, and by pilot awards under ITI Seed 122C158 and CCSB grant U54-CA209971. KB was supported by NIH F32 CA200103; AS by the Mobility Plus fellowship from the Ministry of Science and Higher Education, Poland (1319/MOB/IV/2015/0); SN by the Swedish Society for Medical Research and Stanford Dean’s Fellowship; MR by Swedish Research Council; and HK by an American Heart Association Fellowship.

## Methods

### Mice

Apln^CreER^ (apelin; targeted mutation 1.1, Bin Zhou; kindly provided by Dr. Ralf H Adams), BALB/cJ (The Jackson Laboratory) and B6.129(Cg)-Gt (ROSA)26Sortm4(ACTB-tdTomato,- EGFP)Luo/J (The Jackson Laboratory) mice were bred and maintained in the animal facilities at Veterans Affairs Palo Alto Health Care System, accredited by the Association for Assessment and Accreditation of Laboratory Animal Care. LysM^GFP^ mice (lysozyme 2; targeted mutation 1.1; kindly provided by Dr. Thomas Graf) were maintained in a specific pathogen-free environment at the University of Calgary Animal Resource Centre.

### Preparation of lymphoid tissue BECs for flow cytometry

Axillary, inguinal and brachial PLN from 20-30 adult mice: BALB/cJ (PLN1 and PLN2) or mice of mixed background (PLN3) were dissociated as described^80^. To minimize technical variation, in one study (PLN1) male and female PLN were combined before processing the tissue and separated post-sequencing using the AddModuleScore function from the Seurat package (v3.1.1) to calculated enrichment of male-specific genes (y-chromosomal genes) and the female specific gene *Xist*. Endothelial cells were isolated essentially as described^80^ (online version). Approximately, 5-10×10^4^ BECs (lin^-^Gp38^-^CD31^+^) were sorted into 100% fetal bovine serum using a FACS Aria (100µm nozzle; ∼2500 cells/second). Freshly sorted cell suspensions were diluted with PBS to a final FBS concentration of ∼10% and centrifuged at 400g for 5 minutes. Supernatant was carefully removed using micropipettes and cell pellets resuspended in the residual volume (∼30-50 µl). Cells were counted using a hemocytometer and cell concentration adjusted to 500-1000 cells µl) by addition of PBS with 10% FBS if necessary.

### Single-cell RNA sequencing

Cell suspensions were processed for single-cell RNA-sequencing using Chromium Single Cell 3’ Library and Gel Bead Kit v2 (10X Genomics, PN-120237) according to 10X Genomics guidelines. Libraries were sequenced on an Illumnia NextSeq 500 using 150 cycles high output V2 kit (Read 1-26, Read2-98 and Index 1-8 bases). The Cell Ranger package (v3.0.2) was used to align high quality reads to the mm10 transcriptome (quality control reports available: https://stanford.io/37sXZV3). Normalized log expression values were calculated using the scran package^81^. Imputed expression values were calculated using a customized implementation (https://github.com/kbrulois/magicBatch) of the MAGIC (Markov Affinity-based Graph Imputation of Cells) algorithm^82^ and optimized parameters (t = 2, k = 9, ka = 3).

Highly variable genes were identified using the FindVariableGenes function (Seurat, v2.1) as described^34^. For analyses designed to identify clusters, non-variable genes, cell cycle genes^83^, genes detected in fewer than 3 cells and genes with an average expression level below 0.3 (normalized and log-transformed counts) were excluded. Supervised cell selection was used to remove cells with non-blood endothelial cell gene signatures: lymphatic endothelial cells (*Prox1*, *Lyve1*, *Pdpn*); Pericytes (*Itga7*, *Pdgfrb*); fibroblastic reticular cells (*Pdpn*, *Ccl19*, *Pdgfra*); lymphocytes (*Ptprc*, *Cd52*). Top principal components and the FindClusters function (Seurat, v2.1; res = 0.3) were used on a core set of cells (2394) from the PLN1 sample to identify the 8 major clusters. The Arterial, HEC and CRP clusters were further subdivided into Art and Pre-Art, HEC and HEC (late); and CRP and CRP (early), based on canonical marker expression and their position in tSpace projections of PLN1, yielding a total of 11 subsets. The remaining PLN1 cells and cells from the independently processed samples (PLN2 and PLN3) were assigned the identity of the maximally correlated (Pearson) average expression profile of the core PLN1 cell subsets using ∼3000 common variable genes. Batch effects from technical replicates were removed using the MNN algorithm^84^ as implemented in the batchelor package’s (v1.0.1) fastMNN function. Cells were classified as dividing or resting using a pooled expression value for cell cycle genes (Satija Lab Website: regev_lab_cell_cycle_genes). For UMAP and tSpace embeddings, cell cycle effects were removed by splitting the data into dividing and resting cells and using the fastMNN function to align the dividing cells with their resting counterparts. Dimensionality reduction was performed using the UMAP algorithm (arXiv: 1802.03426) and nearest neighbor alignments for trajectory inference and vascular modeling were calculated using the tSpace algorithm^85^. Cells along isolated trajectories were selected by gating within tPC projections 1-5 as described^85^; and illustrated in the figures here. Differential gene expression analysis was performed by comparing each subset to the remaining cells and fitting a zero-inflated negative binomial model using the LineagePulse package, v0.99.20. In order to assess differentiation potency of single cells, signaling entropy rate (SR) of was calculated as described^46^ after random down-sampling of reads to 1000 reads/cell.

### Data Visualization

Heatmaps were generated using the ComplexHeatmap package^86^, scaled to a maximum value of 1. Data for trajectory heatmaps was pre-processed using code adapted from the plot_as_function from cyt (Pe’er Lab): normalized count data was smoothed with respect to trajectory distance using a gaussian kernel and plotted using a discrete color scale. Violin plots were generated using ggplot2; y-axis units for gene expression data correspond to log-transformed normalized counts after imputation. 3d plots were generated using the rgl package v0.100.30^87^, with minor source code modifications for interactive renderings.

### GO term analysis

Pooled expression values for GO term gene sets and other sets of genes were calculated as previously described^88^ using the AddModuleScore function of Seurat (v3.1.1), which centers values by subtracting pooled expression for random sets of control genes with similar expression levels. To identify biologically relevant GO terms, we first generated a pooled expression matrix by systematically applying the AddModuleScore function to all currently annotated GO terms with at least 3 expressed genes. Differentially regulated GO terms were identified from the resulting “GO term expression matrix” (14300 GO terms by 8832 cells) by comparing each subset to the remaining cells using a Student’s t-Test. A high degree of overlap was observed with conventional GO term analysis approaches such as analysis of top differentially expressed genes using Enrichr. Because this GO term analysis approach is done on a cell by cell basis, it was particularly useful for the identification of terms whose enrichment spanned multiple subsets, e.g. GO:0015669_gas transport (**Supplementary Fig. 3**).

### Data Availability

Data are available from the GEO database (accession GSE140348).

### Antibodies

The following antibodies were used for both microscopy and FACS: Brilliant Violet (BV) 605- conjugated CD31 (390), peridinin chlorophyll protein–cyanine 5.5–conjugated anti-CD45 (30- F11), peridinin chlorophyll protein–cyanine 5.5–conjugated anti-Ter-119 (TER-119), peridinin chlorophyll protein–cyanine 5.5–conjugated anti-CD11a (H155-78), peridinin chlorophyll protein–cyanine 5.5–conjugated anti-CD326 (<u>G8.8</u>), phycoerythrin-cyanine 7-conjugated anti-Gp38 (8.1.1), anti-VE-Cadherin (VECD1), and BV421-conjugated anti-CXCR4 (L276F12) were from Biolegend. BV421-conjugated anti-CD146 (ME-9F1) and BV480 Streptavidin were from BD Biosciences. Anti-estrogen receptor alpha antibody (SP1) and Anti-ERG antibody (EPR3864) were from Abcam. Anti-CD276 (MIH35) and isotype control mouse IgG2a (eBM2a) were from Thermo Fisher Scientific. Anti-PNAd (MECA-79), anti-Ly6c (Monts1), anti-EMCN (5C7), anti-PODXL (MECA-99), anti-ICAM-1 (BE29G1), anti-VCAM-1 (6C2.1), anti-PLVAP (MECA-32) and anti-Slex (F2) were produced in-house from hybridomas; labelled with DyLight Antibody Labeling Kits or Biotin labeling kit (Thermo Fisher Scientific). Alexa Fluor 488–conjugated donkey antibody to rabbit IgG (711-546-152) was from Jackson ImmunoResearch Laboratories. Antibodies for *in vivo* blockade of selectins were from BD Biosciences: Anti-E-selectin (10E9.6), anti-P-selectin (RB40.34); and eBioscince: isotype control rat IgG1 (NALE).

### Imaging

PLN were imaged following either retroorbital injection of fluorescent labeled antibodies or by fluorescence staining of LN sections. If injected, antibodies (25-75 µg) were administered 5-30 minutes prior to sacrifice and PLN removal. To image the overall vascular, the PLN was gently compressed to ∼35-50 µm thickness on a glass slide. Alternatively, PLNs were fixed with 4% paraformaldehyde, cryoprotected with sucrose, frozen in OCT (Sakura® Finetek) in 2- methylbutane (Sigma) on dry ice and stored at −20 °C. 50 µm cryo-sections were stained with antibodies according to standard protocols. The slides were imaged using Apotome 2.0 fluorescence microscope or LSM 880 laser scanning microscope (Zeiss).

For quantification of ER^+^ cells, anti-human estrogen receptor antibody was used as a surrogate stain for Apln in sections from Apln^CreER^ mice. Capillaries, HEV and ER^+^ vessels were enumerated within 50 µm sections at 20X objective using a grid reticle to determine the relative frequency of each EC subset (1 length unit = ⅛ of the grid height). Sufficient fields were scanned to comprise >5000 HEC assessed for reactivity with anti-ER antibody. Data was expressed as frequency of ER^+^ (as a % of total counted vessels, i.e., length of ER^+^/ length of capillaries + HEC) per lymph node section. We scanned nine LN from two Apln^CreER^ unchallenged mice and nine LN from two Apln^CreER^ mice five days post cutaneous inflammation (one section per LN). The number of HEC per field was determined based on ERG^+^ nuclei within MECA79^+^ EC using 10X or 20X objectives.

### Live Imaging

LysM^GFP^ mice were injected in the right footpad with 2.5 × 10^7^ CFU *S. aureus.* An hour later mice received intravenously injected Cell Tracker Red CMTPX (Thermofisher) labeled lymphocytes. An hour after lymphocyte injection mice were injected intravenously with a mixture of anti-PNAd- Dylight594 and albumin-Dylight680 to visualize HEV and vasculature, respectively. Two-photon video microscopy was employed to assess the movement and location of neutrophils and lymphocytes in lymph node blood vessels following infection with *S. aureus.* Right hindlimb popliteal LN was exposed for imaging in the anesthetized mouse. The LN was imaged from 2 to 4 hpi. Image acquisition was performed an upright two-photon microscope (Leica Biosystems TCS SP8 Upright Microscope). For antibody blocking studies, mice were pretreated with blocking antibody 20 minutes before infection.

### Lymph Node Immunization

Oxazolone: Mice were subjected to cutaneous immune challenge by applying 20 µl of 3-5% 4- Ethoxymethylene-2-phenyl-2-oxazolin-5-one (Sigma-Aldrich) in 1:2 acetone:olive oil. Peripheral LN (axillary, brachial, inguinal) were harvested at varying timepoints post inflammation for imaging. Complete Freund’s adjuvant (CFA): Mice received a unilateral hock injection of 10 µl Complete Freund’s Adjuvant (Sigma-Aldrich), the popliteal LN were harvested three and half weeks later, and the inflamed nodes were compared to uninflamed control nodes with imaging.

### Lineage tracing

For lineage tracing, reporter expression was induced in Apln-CreER x R26-mTmG by i.p. injection of 80 microg/g of 4-hydroxytamoxifen (4-OHT; Sigma-Aldrich) 24 hours prior to sacrifice, or 76 hours prior to sacrifice in separate experiments, for imaging or to the start of immunization. Reporter expression was induced in Apln CreER x R26-tdTomato mice the day after oxazolone skin painting, and lymph nodes were imaged 24 hours later or after 11 days.

### Statistical Analysis

Statistical significance between two groups was calculated using a two-way ANOVA corrected with Tukey. The rule of three was applied to determine 95% confidence intervals for the enumeration of ER^+^ cells. A likelihood ratio test was used for differential gene expression analysis assuming an underlying zero-inflated negative binomial distribution. P values were adjusted for multiple comparisons by calculating the false discovery rate (FDR) and adjusted p-values < 0.001 were considered significant.

---

**Supplementary Figure 1.**
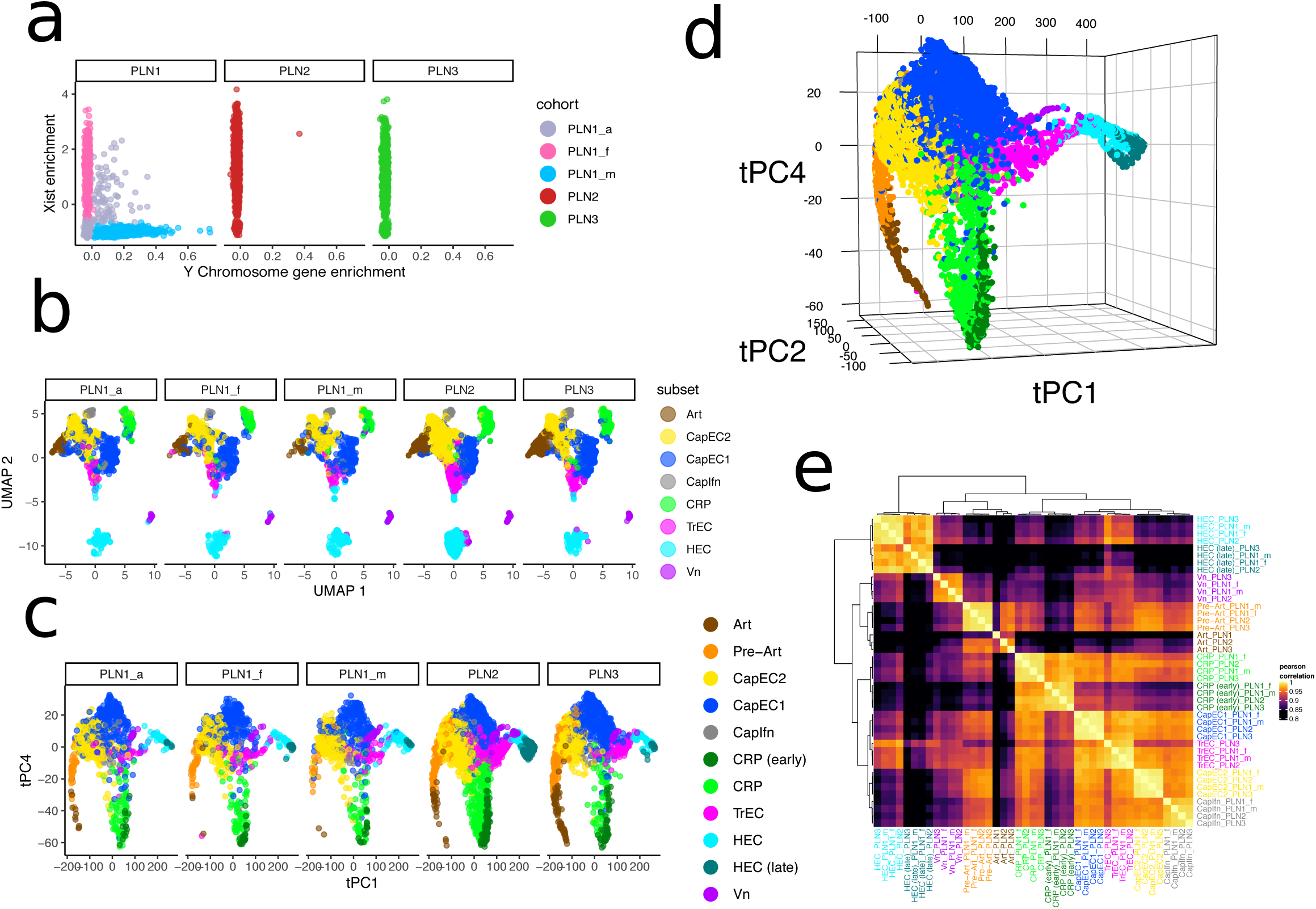
Consistency of three technical replicates and four independent mouse cohorts. (a) Scatter plots of all cells showing separation of the PLN1 sample into male and female cohorts (PLN1_m and PLN_f) and remaining unclassifiable PLN1_a cells. (b) UMAP plot from Fig. 1d, stratified by cohort. (c) and (d) tSpace projection (2D in (c) and 3D in (d)) using all cells colored by subset. Interactive rendering available: https://stanford.io/2WXR811 (e) Pearson correlation of gene expression profiles of subsets from different cohorts. A set of ∼2000 differentially expressed genes was used to calculate mean expression profiles for each of the major subsets (Art, CapEC2, CapEC1, CapIfn, CRP, TrEC, HEC, Vn) in each cohort (PLN1_m, PLN1_f, PLN2 and PLN3). Cells of the Art subset from the PLN1_m and PLN_f cohorts were combined and treated as a single cohort due to low total number of Art cells in PLN1_m. Expression profiles were hierarchically clustered and plotted along with their pairwise pearson correlation coefficients (color scale).

**Supplementary Figure 2.**
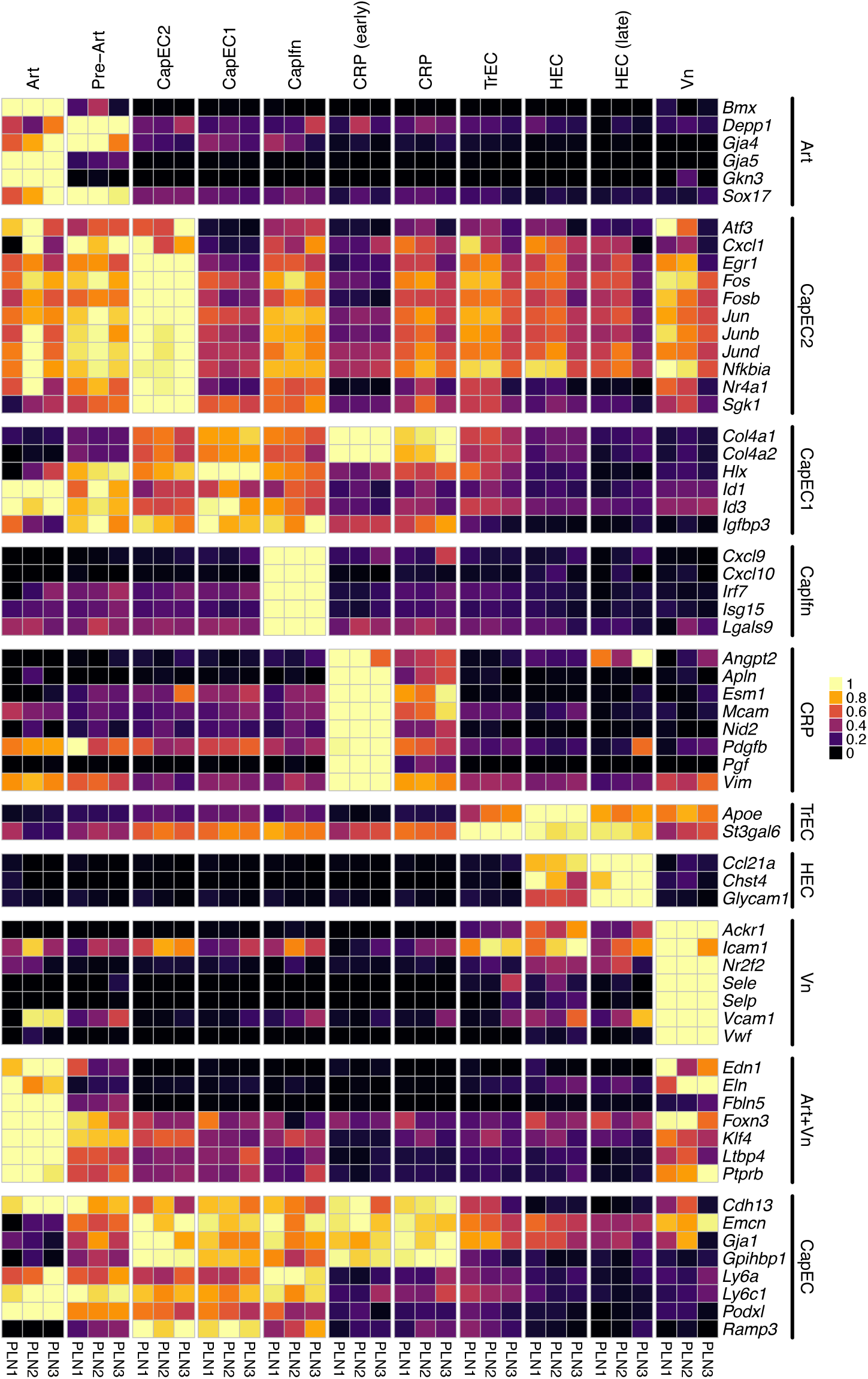
Consistency of gene signatures from. Fig 1 **across 3 replicates.** Average expression of genes shown in Figure 1f. Scaled from 0 to max on a per sample basis.

**Supplementary Figure 3.**
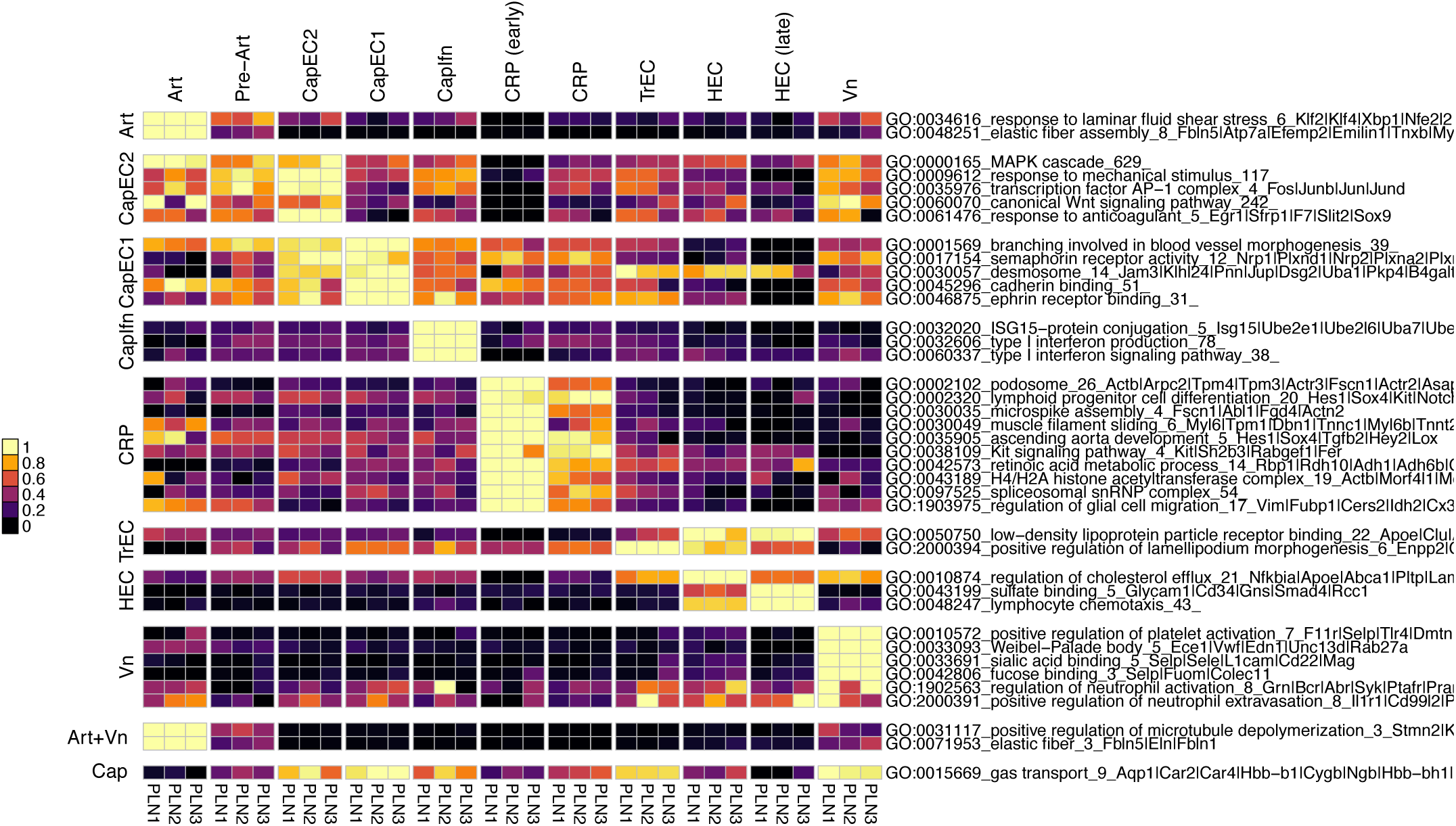
Pooled expression of genes grouped according to their Gene Ontology (GO terms). Average pooled expression for select differentially enriched GO terms in each sample. The number of genes belonging to a given term is indicated after the term identifiers. When feasible, symbols for individual genes (separated by a “|”) belonging to given GO term are listed in order of highest to lowest expression across all datasets.

**Supplementary Figure 4.**
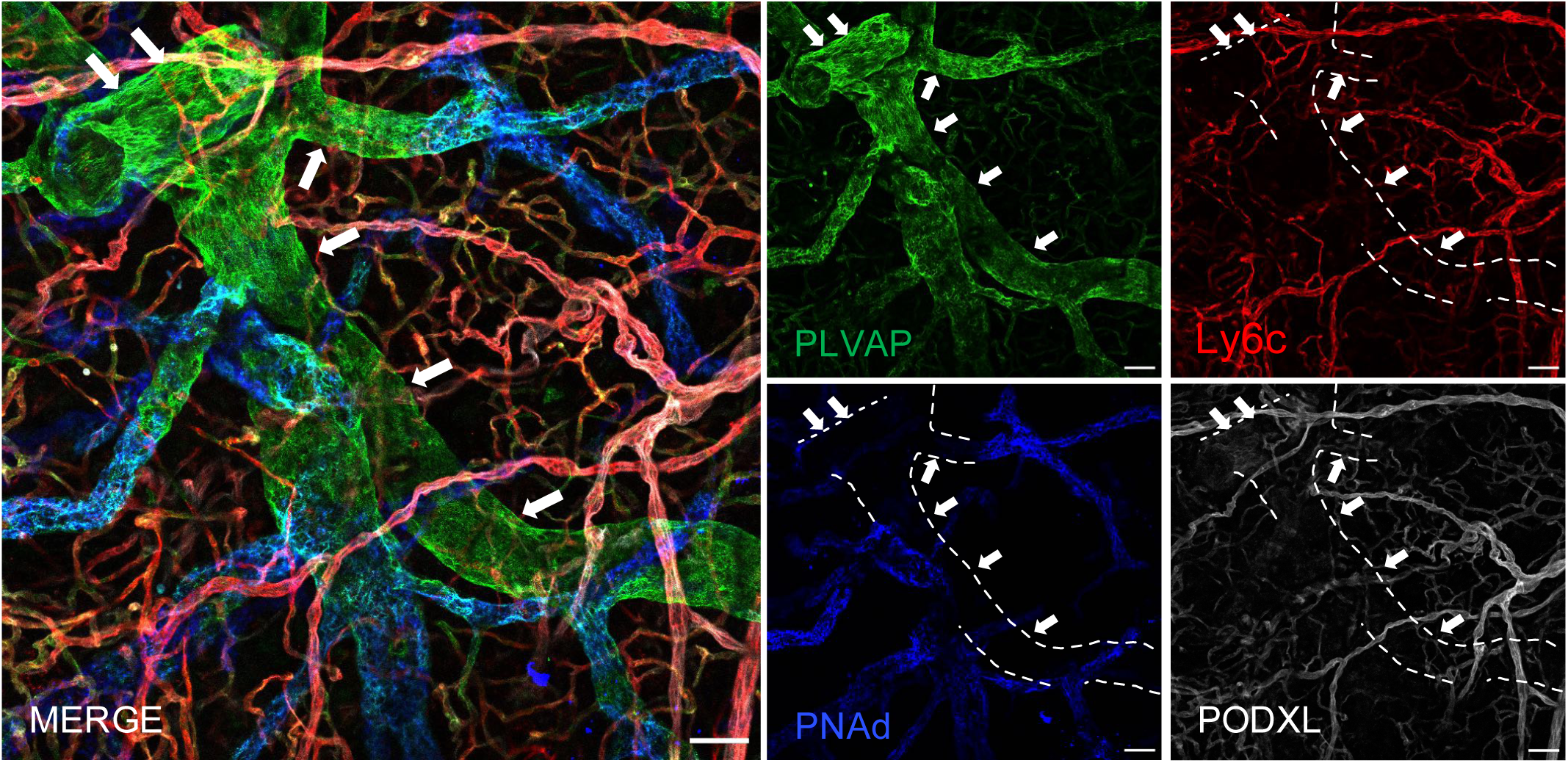
Additional image illustrating the medullary vein subset. Immunofluorescent image of PLN stained with i.v. injected anti-PLVAP (green), anti-Ly6c (red), anti-PNAd (blue) and anti-PODXL (white). Scale bar 50 μm. Arrows point to medullary vein.

**Supplementary Figure 5.**
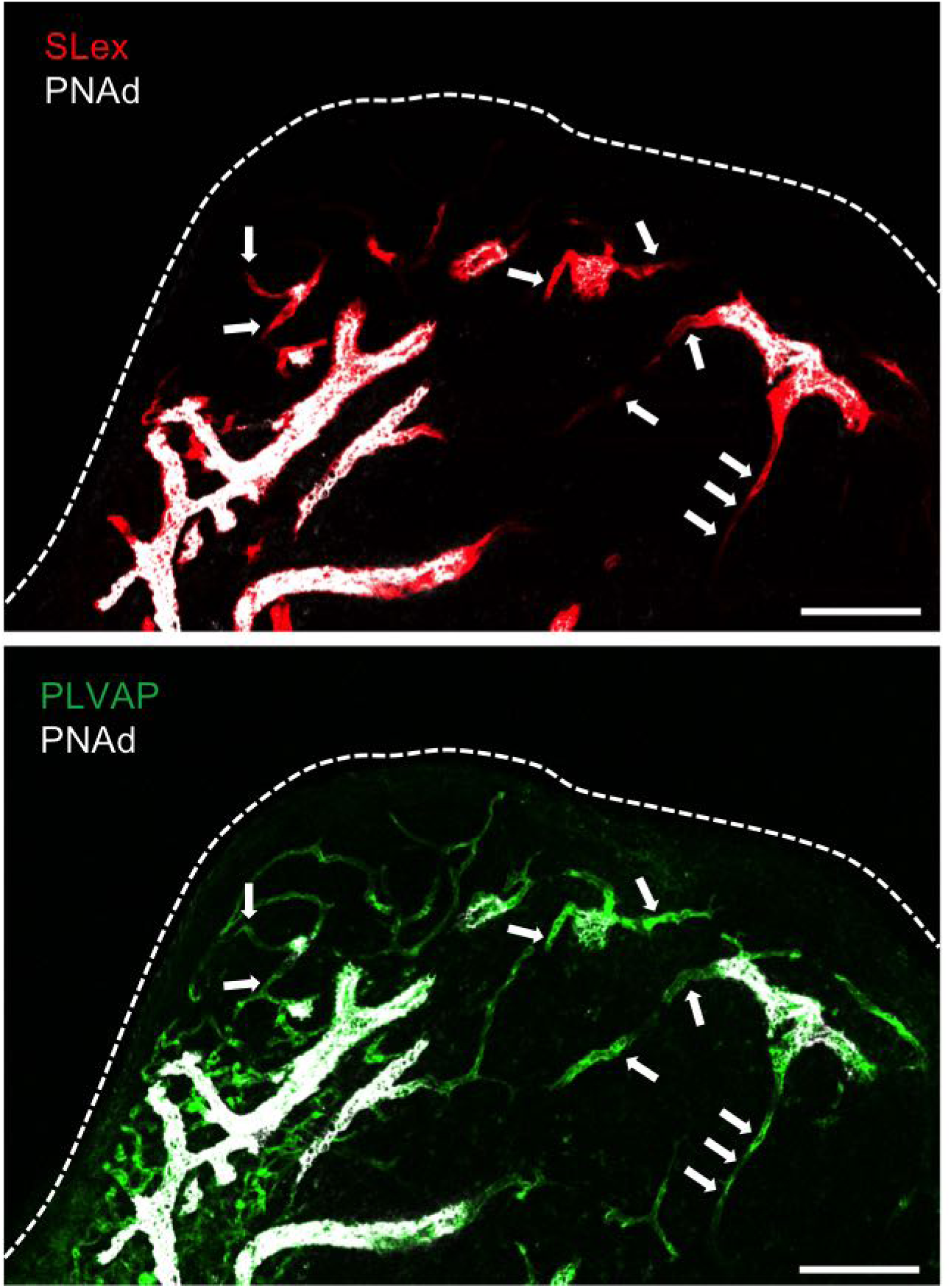
TrEC localize to capillary segments entering HEV. Immunofluorescent image of PLN stained with i.v. injected anti-sLex (red), anti-PNAd (white) and anti-PLVAP (green). Arrows indicate TrEC, capillary segments expressing sLex but not PNAd . Bars, 100 µm. Dashed line, LN capsule.

**Supplementary Figure 6.**
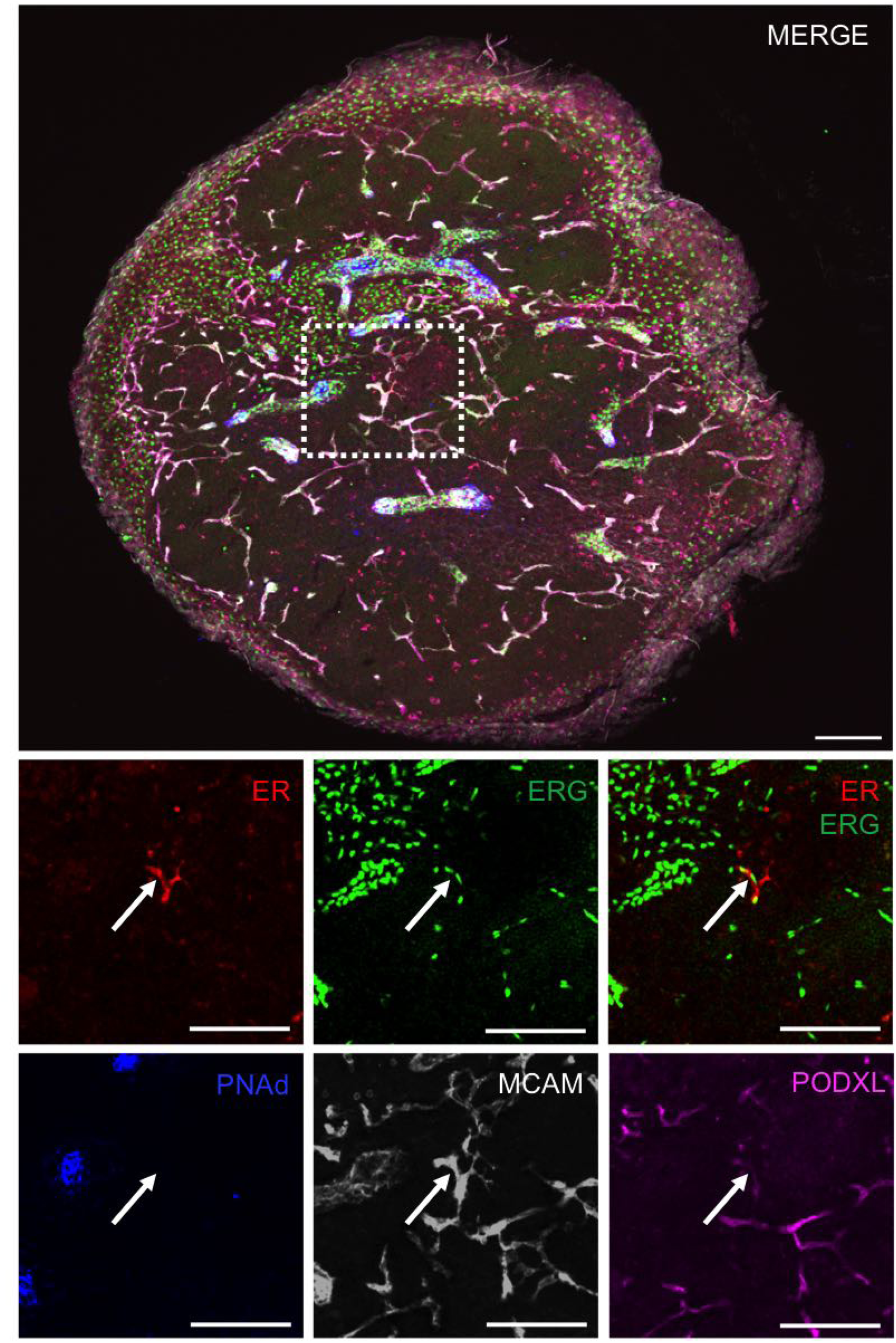
Additional markers characterizing the ER+ CRP subset. Immunofluorescent image of PLN stained with anti-ER (red), anti-ERG (green), anti-PNAd (blue), anti-MCAM (white) and anti-PODXL (violet). Scale bar 100 µm. Arrows point to CRP.

**Supplementary Figure 7.**
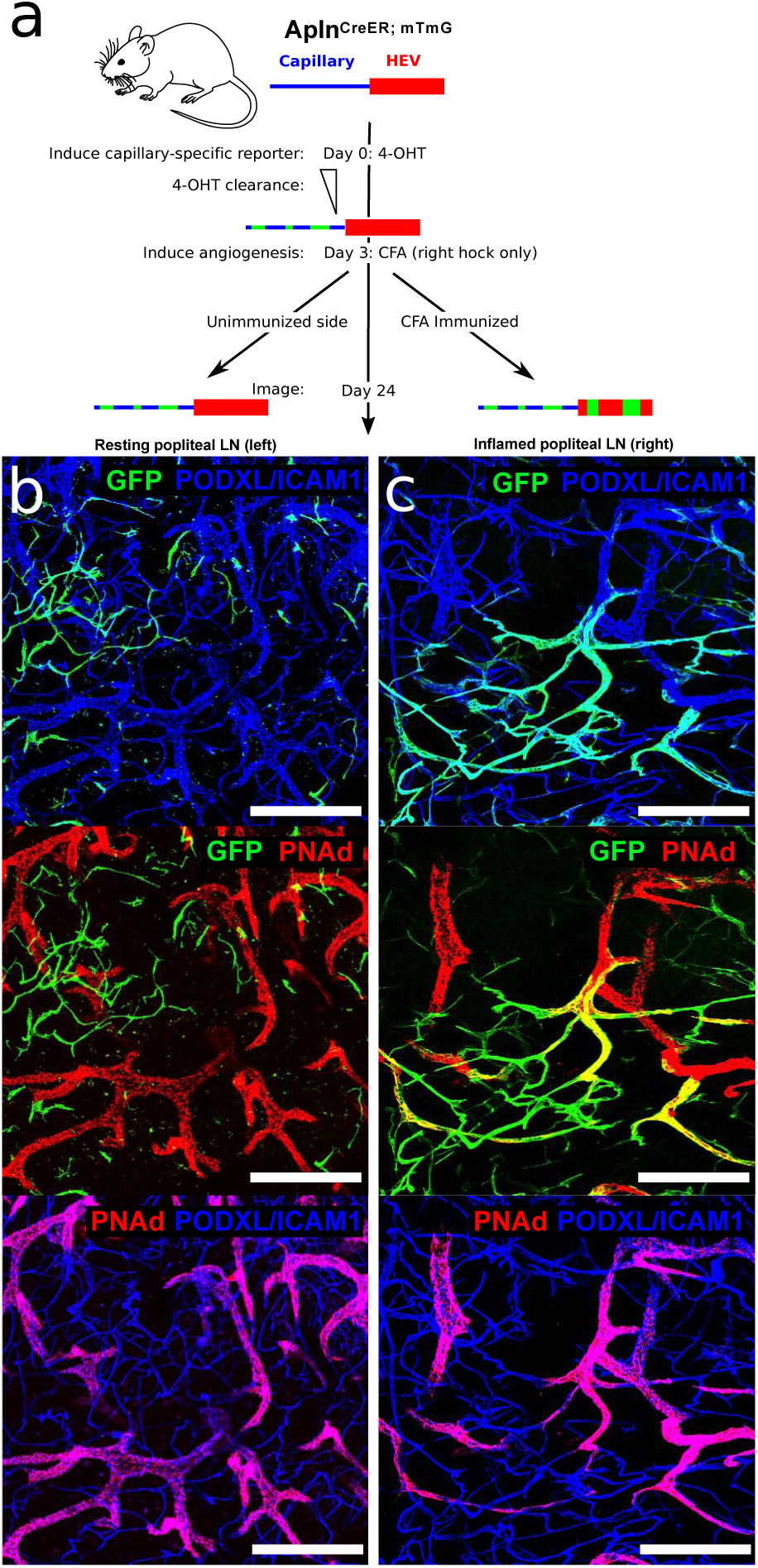
Lineage tracing of AplnERTCre-expressing capillary EC with additional 4-OHT clearance time. (a) Experimental timeline for (b) and (c). Reporter expression was induced in Apln-CreER-mTmG mice by i.p. injection of 4-OHT. 72 hours later CFA was injected into the right hock and three and half week later mice were sacrificed. EC subsets were labeled by i.v. injection of the indicated antibodies 10-20 minutes before sacrifice.: anti-PNAd (red), anti-PODXL (blue) and anti-ICAM1 (blue). Representative images of resting (b) and inflamed (c) popliteal lymph nodes. False color used to represent fluorophores. tdTomato not shown. Bars, 200 µm.

**Supplementary Figure 8.**
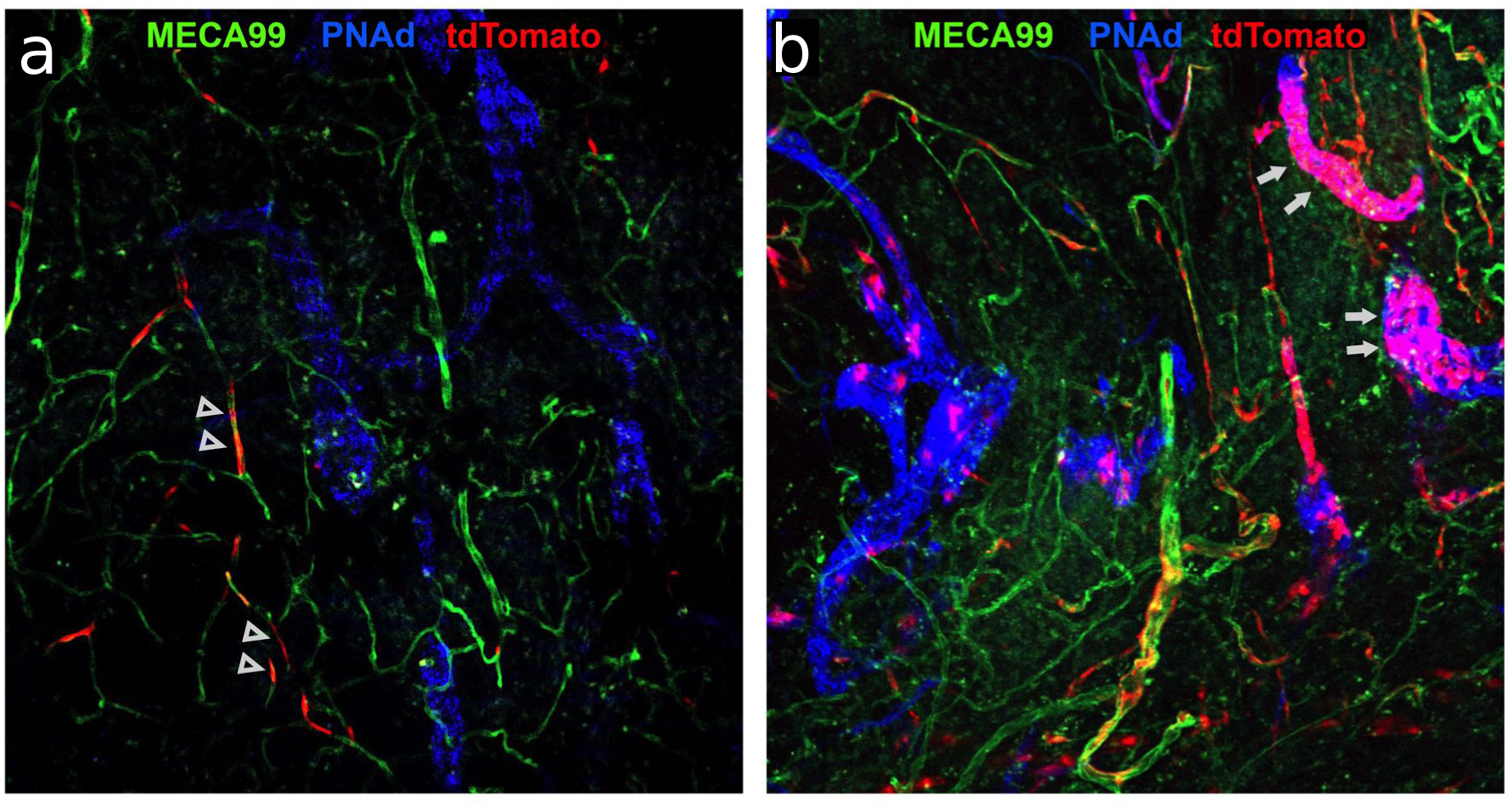
Tamoxifen administration during the early oxazolone response: selective capillary reporter induction and lineage tracing Apln-CreERT2-tdTomato mice were immunized by cutaneous application of oxazolone, pulsed with i.p. tamoxifen the next day, and sacrificed either 48 hours (a) or twelve days (b) after immunization. EC subsets were labeled by i.v. injection of the indicated antibodies 10-20 minutes before sacrifice, and draining lymph nodes were imaged: PODXL (MECA99; green), PNAd (blue), tdTomato (red). Representative images of reporter (tdTomato) positive EC 48 hours (left) or twelve days (right) after immunization. Scale bar represents 50 μm.

**Supplementary Figure 9.**
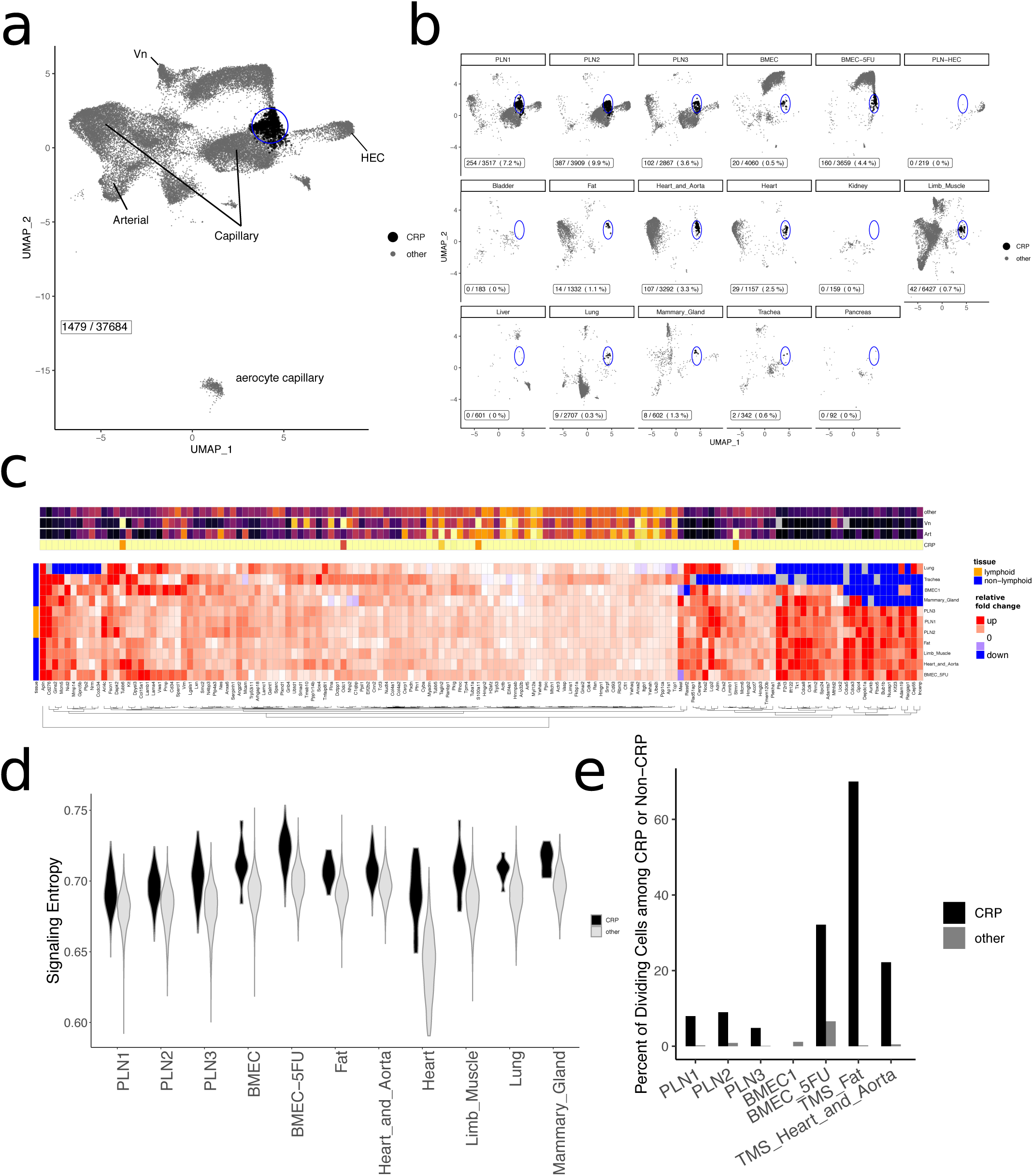
Global alignment of PLN BEC with publicly available single cell BEC profiles. (a and b) UMAP plot of MNN-aligned data from various public BEC datasets. a) all BEC aligned, illustrating the position of CRP-like EC. b) Separate plots for each tissue. Blue circle highlights the region to which LN CRP map. Black dots represent cells that align with LN CRP and that also are mare similar in gene expression profile to CRP than to other LN EC subsets. PLN samples are from this study. PLN-HEC are sorted EC from mice55. BMEC1 and BMEC_5FU are sorted Cdh5-reporter-positive EC from bone marrow54. Other samples are from Tabula Muris consortium53. (c) Heat maps of genes selectively expressed by LN CRP and CRP-like EC in other tissues. Top panel shows selectivity compared with other EC subsets (combined data all tissues). Bottom panel shows fold-change analysis of CRP-like EC compared to non-CRP in different tissues, and illustrates shared expression. (d) Violin plots of the signaling entropy rate of the indicated samples. (e) Bar graph depicting the percentage of CRP or of non-CRP that express signatures of cell division in each of the indicated samples.

**Supplementary Figure 10.**
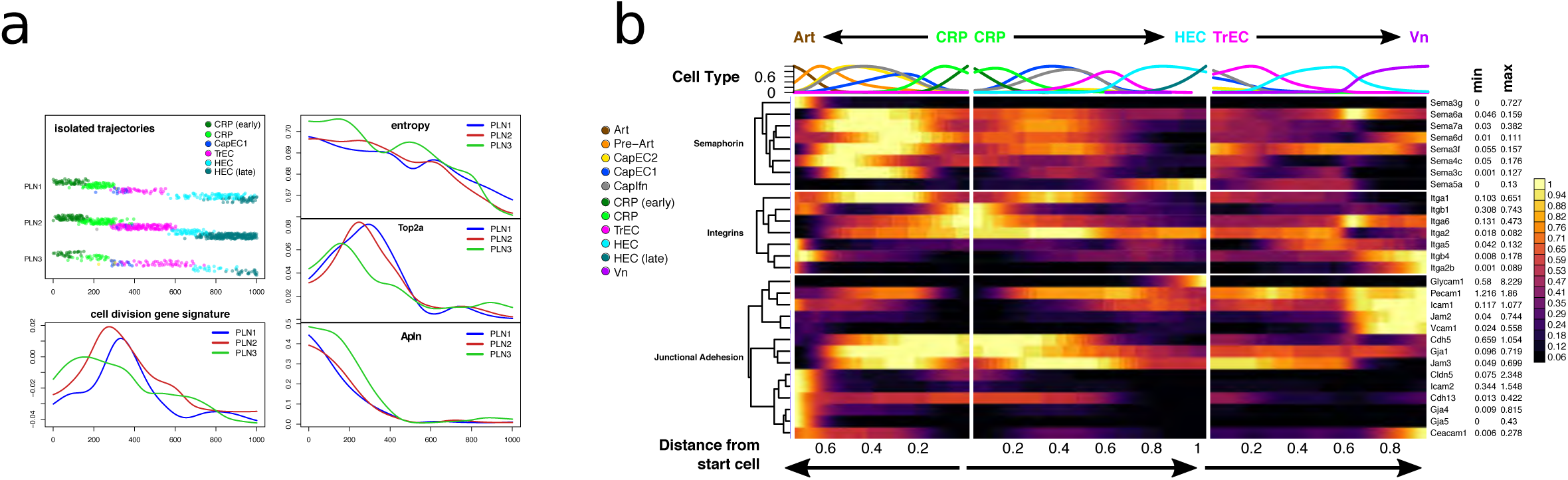
Expression of additional genes and features along isolated trajectories. (a) Expression of select genes and features plotted along cell trajectories from early CRP to HEC and smoothed as in Figure 7. Apln is downregulated rapidly from early to late CRP, while markers of cell cycle (e.g. Top2a, and a global division signature), increase and peak in late CRP and TrEC. The “cell division gene signature” was quantified as a pooled expression value of previously defined cell division genes83. (b) Expression of selected genes along cell trajectories from early CRP to Art (plotted leftward), and from CRP to HEC or Vn (rightward) as in Figure 7.

